# *In situ* distance measurements in a membrane transporter using maleimide functionalized orthogonal spin labels and 5-pulse electron double resonance spectroscopy

**DOI:** 10.1101/2021.12.23.473964

**Authors:** Sophie Ketter, Marina Dajka, Olga Rogozhnikova, Sergey A. Dobrynin, Victor M. Tormyshev, Elena G. Bagryanskaya, Benesh Joseph

## Abstract

Spectroscopic investigation of membrane proteins in their native environment is a challenging task. Earlier we demonstrated the feasibility to measure precise distances within outer membrane proteins in *E. coli* and native membranes using methanethiosulfonate (MTS) functionalized labels combined with pulsed electron double resonance spectroscopy. Here we show the application of maleimide functionalized Gd(III), nitroxide, and trityl labels for *in situ* distance measurement using the cobalamin transporter BtuB. These labels enabled distance measurements for BtuB in *E. coli* and native outer membranes and in the membranes maleimide-Gd-DOTA also is effective. Further, we show that the observable dipolar evolution time can be significantly prolonged in the native environments using the Carr-Purcell 5-pulse electron double resonance sequence. For a nitroxide-nitroxide pair, application of sech/tanh inversion pulses substantially suppressed the 4-pulse artifact at the Q-band frequency. In the case of a nitroxide-trityl pair, Gaussian pump pulses of varying amplitude are sufficient to suppress the artifact to the typical noise level. The feasibility of a range of bioresistant spin labels and the 5-pulse electron double resonance offers promising tools for investigating heterooligomeric membrane protein complexes in their native environment.

## 1. Introduction

Pulsed dipolar electron spin resonance spectroscopy techniques are widely used to measure the dynamics of biomolecules. Double electron-electron resonance (DEER or PELDOR) is the most popular among these techniques. The initial 3-pulse sequence, which was used to observe the dipolar coupling between nitroxide spins was later extended into a deadtime-free 4-pulse sequence [1, 2]. Since then, it has become the standard technique for measuring long-range distances, especially for proteins and nucleic acids [3-9]. In the 4-pulse PELDOR experiment, the observable dipolar evolution time (and the measurable distance) is limited by the phase memory time (*T*_*M*_) of the spins. For protein molecules in aqueous solution, *T*_*M*_ is dominated by nuclear spin diffusion [10-12]. In such cases, deuteriation can significantly prolong the *T*_*M*_ and the accessible distances [13, 14]. The *T*_*M*_ is considerably reduced in the membrane environment (1-2 µs), where the higher local concentration or the rotating methyl groups in proteins and lipids were suggested to make a major contribution [15, 16]. Here, enhancing the *T*_*M*_ with a complete deuteriation of the protein and the surrounding lipids is not always a feasible strategy.

To overcome the effect of this diffusive process on the phase memory of electron spins, Borbat *et al*., introduced a 5-pulse sequence [17]. Here, the observer pulses sequence is similar to the 4-pulse PELDOR, but with a symmetrized placing as in a Carr-Purcell sequence. An additional static pump pulse is applied between the first two observer pulses, which shifts the phase of the coupled spins such that the dipolar evolution can be observed for the entire time between the last two observer pulses. Due to the non-uniform inversion probability of the successive pump pulses, a partial excitation (4-pulse) artifact is overlaid on the primary data. The authors suggested that this artifact could be reduced with a stronger standing pump pulse or even eliminated with broad-band shaped pulses having a uniform population inversion. Over the years, research from the Jeschke and the Prisner groups led to the implementation of an Arbitrary Waveform Generator (AWG) for applying such shaped pulses [18-21]. Among these pulses, the hyperbolic secant/hyperbolic tangent (sech/tanh) pulses were shown to provide the highest selectivity and asymmetric pulses of the same length and bandwidth achieve even higher adiabaticity [22, 23]. At the X-band (9.4 GHz) frequency, such asymmetric sech/tanh pulses suppressed the partial excitation artifact below the noise level [24]. However, similar performance could not be observed at the Q-band (34 GHz) frequency, at which most of the PELDOR experiments are routinely performed.

In the past decade, several studies on the application of PELDOR for measuring distances within the native environment of the biomolecules have been reported [25-33]. This was facilitated through the developments in pulsed ESR spectrometer instrumentation, redox stable spin labels, and new labeling strategies and methods for delivering labeled molecules into the cells [18, 19, 26, 34-41]. For soluble proteins and nucleic acids, pre-labeled molecules can be introduced into the cells [26, 40, 42, 43]. This approach is not feasible for membrane proteins, and direct *in situ* labeling approaches are necessary. Labeling in the complex membrane environment poses many difficulties. With the technologies available today for bacterial expression, it is feasible to express the target membrane protein to the desired level for ESR spectroscopy (∼10^5^ copies/cell). Nonetheless, selective labeling of the target site(s) as well as stability of the spin label in the complex environment are the major challenges. Further, accessibility of the target sites could be considerably limited.

Over the past years, we demonstrated the feasibility to perform distance measurements on outer membrane proteins (OMPs) of Gram-negative bacteria in the native outer membranes (OM) and intact cells [39, 44-46]. The cobalamin transporter BtuB and the protein insertase BamA were studied in the native environment using this approach [46-48]. The outer membrane (OM) is an asymmetric bilayer consisting of phospholipids and lipopolysaccharides (LPS). It harbors numerous β-barrel proteins performing crucial functions including LPS transport and OMP folding. In *E. coli*, OMPs rarely have reactive cysteines, which enabled specific labeling of engineered cysteines using the methanethiosulfonate spin label (MTSL). Application of this approach with isolated outer membranes produced more background labeling; nevertheless, a random distribution of these signals does not interfere with distance measurements, albeit somewhat lowering the effective modulation depth [31, 46]. So far, we employed thiosulfonate functionalized nitroxide (MTSL) or trityl (TAM or OX063) labels for these experiments [45, 49, 50]. A combination of nitroxide (NO) and the trityl labels provided higher sensitivity enabling distance determination in BtuB at low micro molar concentrations in the native environments [50]. However, earlier we showed that *E. coli* actively reduces MTSL in a cell density dependent manner [50]. Also, the S-S bond linking these labels with the protein could easily be reduced in the cellular environment and the covalent C-S bond formed by maleimide is more stable [26]. In this study we investigated the feasibility of different maleimide functionalized and bioresistant labels for spin labeling of BtuB. We demonstrate the application of maleimide functionalized nitroxide (3-maleimido-PROXYL (M-PROXYL)) and a *gem*-diethyl nitroxide (MAG1)), trityl (M-OX063), and Gd(III) (M-Gd-DOTA) labels for distance measurements in *E. coli* and native OM. Further, the application of the Carr-Purcell 5-pulse PELDOR sequence for determining long-range distances for membrane proteins within the micellar and the native environments is demonstrated. We show that the sech/tanh pulses can almost completely suppress the 4-pulse artifact for nitroxide labels at the Q-band frequency. For the trityl-nitroxide case, Gaussian pump pulses of different amplitudes are sufficient to suppress the artifact to the typical noise level observed in PELDOR spectroscopy.

## 2. Materials and Methods

### 2.1 Site-directed mutagenesis and protein expression

The cysteine variants of BtuB were created using site-directed mutagenesis as described earlier [44]. The pAG1 plasmid encoding the wild-type (WT) protein or the cysteine variants V90C, T188C, F404C, T426C was transformed into *E. coli* strains RK5016 (*metE70 argH btuB recA*) or BL21(DE3). Cells were grown at 33°C in minimal medium consisting of 0.6 M K_2_HPO_4_, 0.33 M KH_2_PO_4_, 0.08 M (NH_4_)_2_SO_4_, 0.02 M sodium citrate supplemented with 0.01 % (w/v) L-arginine and L-methionine, 0.2 % (w/v) glucose, 150 µM thiamine, 3 mM MgSO_4_, 300 µM CaCl_2_, and 100 µg/mL ampicillin, until an OD_600_ of 0.3–0.4. The cells were diluted 1:650 times into fresh minimal medium and grown overnight at 33°C until an OD_600_ of 0.4-0.5. To perform spin labeling in *E. coli*, the culture was subsequently transferred to ice. To isolate the outer membrane, cells were harvested by centrifugation at 12,000x*g*. LptB_2_FG was expressed from a pETDuet-1 vector following a protocol we described earlier for the related protein TmrAB [51].

### 2.2 Isolation of native outer membranes

Isolation of the outer membrane was done as described earlier [31]. Briefly, the total membrane (∼0.28 g) was suspended into a buffer containing 50 mM MOPS pH 7.5 and 60 mM NaCl followed by a treatment with 0.5 % N-Lauroylsarcosine sodium salt to solubilize the inner membrane. The outer membrane fraction (∼0.1 g) was collected by ultracentrifugation at 200,000x*g* for 1.5 h and suspended in 10 mL of MOPS–NaCl buffer. The pH of the buffer was adjusted to 7.5 for spin labeling with methanethiosulfonate functionalized labels or to 6.8 for maleimide functionalized spin labels.

### 2.3 Preparation of M-Gd-DOTA spin label

The maleimide-Gd-DOTA spin label was prepared by dissolving Maleimido-mono-amide-DOTA (Macrocyclics) and Gadolinium(III) chloride (Sigma Aldrich) in a ratio of 1:1.1 (by mol) in Milli-Q water. The mixture was stirred at room temperature for 8 h while the pH was slowly adjusted to 5.5– 6 using NaOH and further incubated at room temperature overnight. The mixture was then freeze-dried and the resulting powder was dissolved in anhydrous N,N-Dimethylformamide (Sigma Aldrich) and stored under inert atmosphere.

### 2.4 Synthesis of maleimide OX063 (M-OX063) and maleimide-functionalized *gem*-diethyl nitroxide (MAG1)

OX063 (125 mg, 0.092 mmol, 1 equivalent) was prepared as previously reported [52, 53]. 1-hydroxybenzotriazole monohydrate (HOBt, 20 mg, 0.128 mmol, 1.4 equivalent), dicyclohexylcarbodiimide (DCC, 23 mg, 0.110 mmol, 1.2 equivalent) and N,N-diisopropylethylamine (DIPEA, 36 mg, 0.275 mmol, 3 equivalent) were dissolved in anhydrous dimethylformamide (1.5 mL) and stirred under argon at room temperature for 2 h. Hydrochloride of N-(2-aminoethyl)maleimide (23 mg, 0.128 mmol, 1.4 eq) was added to give a deep-green solution, which was stirred under argon overnight. The mixture was transferred into 50 mL beaker and diluted with water (10 mL). To remove solid contaminations, the resulting turbid solution was passed through a short cotton plug. The filtrate was acidified with 0.2 M aqueous HCl to pH 2-3 and concentrated in vacuo to give a brown cake. The product was isolated by column chromatography (silica gel, DCM/methanol from 4:1 to 1:2). This purification procedure was repeated one more time to give the trityl M-OX063 (fine deep-brown powder, 46 mg, 34 %, Figures S17-S18). Data for M-OX063: m.p. > 230 °C (decomposition). HRMS (MALDI TOF): calculated for C_58_H_69_N_2_NaO_19_S_12_ [M+Na]+ 1504,1041, found 1504.213. IR (KBr): ν (cm-1) = 2923 (s), 2881 (s), 1707 (s), 1637 (s), 1581 (vs), 1493 (m), 1437 (s), 1375 (s), 1350 (s), 1313 (s), 1240 (vs), 1039 (vs), 723 (m), 696(m), 669 (m), 660 (m), 636 (m). The maleimide-functionalized gem-diethyl nitroxide (MAG1) was synthesized as described recently [54].

### 2.5 Spin labeling of BtuB in *E. coli*

Following overnight expression, cells were collected by centrifugation at 6,000x*g* for 10 min and suspended in 50 mM MOPS, 60 mM NaCl buffer to reach an OD_600_ value of 0.5 (40 mL total volume). The pH of the buffer was adjusted to the desired spin label as described above. The cells expressing either BtuB WT or a cysteine variant were incubated with 15 µM of either 1-oxyl2,2,5,5-tetramethyl-3-pyrroline-3-methyl methanethiosulfonate (MTSL, Toronto Research Chemicals), 3-Maleimido-2,2,5,5-tetramethyl-1-pyrrolidinyloxy (M-PROXYL, Sigma Aldrich), 3-((2,5-dioxo-2,5-dihydro-1*H*-pyrrol-1-yl)methyl)-2,2,5,5-tetraethylpyrrolidin-1-oxyl (MAG1), maleimide-OX063 (M-OX063), or maleimide-Gd-DOTA (M-Gd-DOTA) for 1 h at room temperature. Excess spin label was removed by three rounds of centrifugation at 6,000x*g* and gentle but thorough suspension in the same buffer. Finally, the cells were suspended in 35–45 µL and transferred to a micropipette (0.64-, 0.86-, or 1.2 mm inner diameter, BLAUBRAND, intraMARK) for the ESR experiment.

### 2.6 Spin labeling and quantification of BtuB in the native outer membranes

Spin labeling was performed with 10 µM final concentration for 1 h at room temperature for all the spin labels. In the case of the periplasmic cysteine variant T426C, 25 µM of the methanethiosulfonate functionalized OX063 (MTS-OX063) spin label was introduced by five cycles of freeze-thawing of the outer membrane. The membranes were subjected to three rounds of ultracentrifugation at 200,000xg for 1.5 h in 60 mL to remove unbound spin label, which always included five cycles of freeze-thawing at each round for the T426C mutant. The final outer membrane pellet was suspended in 60–90 µl of buffer. A two-to three-fold dilution of the sample was transferred to a micropipette (0.86-or 1.2 mm inner diameter) for the cw ESR measurement. The amount of BtuB was assessed by a titration of the TEMPO labeled cyanocobalamin substrate (T-CNCbl) as we previously described [44].

### 2.7 Sample preparation for the kinetic analysis of spin label stability

Following overnight expression, cells overexpressing BtuB WT protein were pelleted at 6,000x*g* and suspended into an appropriate volume of 50 mM MOPS pH 7.5 and 60 mM NaCl to reach the OD_600_ values of 0.5 or 5. The spin label was added to a final concentration of 200 µM, mixed thoroughly but gentle, and 50 µL were transferred to a 1.2 mm inner diameter micropipette. For the control experiments in buffer or to assess the effect of ascorbic acid, spin labels were added to 200 µM final concentration into the same buffer containing 1- or 5 mM L-ascorbic acid (Sigma Aldrich).

### 2.8 Continuous-wave ESR spectroscopy

All continuous-wave (cw) ESR spectra were recorded at room temperature as the first derivative signal on a Bruker EMXnano Benchtop Spectrometer operating at X-band frequency (9.4 GHz). Nitroxide spectra of spin labeled samples were acquired with 100 kHz modulation frequency, 0.15 mT modulation amplitude, 2.0 mW microwave power, 5.12 ms time constant, 22.5 ms conversion time and 18 mT sweep width. For trityl labeled samples, spectra were recorded with 100 kHz modulation frequency, 0.02 mT modulation amplitude, 1.0–2.0 mW microwave power, 1.28 ms time constant, 3–4.8 ms conversion time, and 5 mT sweep width. For whole cell samples, spectra were averaged over 40 scans with a sweep time of 27 s for nitroxide and over 150–350 scans with a sweep time of 7.5–12 s for trityl. For outer membrane samples, spectra were averaged over 12– 156 scans with sweep times as given above. The spin concentration of the samples was quantified using the reference free concentration determination of paramagnetic species as implemented in the Xenon software (Bruker). For quantitative comparison, the cw spectra of spin labeled BtuB in *E. coli* were normalized to a same OD_600_ value. For the outer membrane samples, the cw spectra were normalized to the same amount of BtuB (as quantified using T-CNCbl) unless otherwise stated.

The parameters for the kinetic analysis of nitroxide spin labels in *E. coli* suspension were 100 kHz modulation frequency, 0.1 mT modulation amplitude, 2.0 mW microwave power, 5.12 ms time constant, 12–15 ms conversion time and 10 mT sweep width. Spectra were recorded with a delay of 30 s and averaged over 4 scans with sweep time of 15 s. For trityl in *E. coli* suspension, a 100 kHz modulation frequency, 0.02 mT modulation amplitude, 0.6–2.0 mW microwave power, 1.28 ms time constant, 4–5 ms conversion time, and a 2.5 mT sweep width was used. The delay was set to 10 s and spectra were averaged over 2 scans with a sweep time of 5 s. For the measurements in buffer or in the presence of ascorbic acid, the delay was set to 60 s for nitroxide and trityl spectra were recorded with a delay of 40 s and averaged over 8 scans. Data points were extracted from the double integral of the spectra after taking the delay for the start of the measurement into account (2–3 min).

### 2.9 Pulsed ESR spectroscopy

All the experiments were performed on a Bruker Elexsys E580 Q-band Pulsed ESR spectrometer with SpinJet AWG, which was recently installed in our work group. It is equipped with a continuous-flow helium cryostat, a temperature control system (Oxford Instruments), a 50 W solid state amplifier, and a Bruker EN5107D2 dielectric resonator. The samples contained 20 % of deuterated glycerol (v/v) as well as T-CNCbl when required. For *E. coli* samples, ∼20–30 µM T-CNCbl was added to all the samples, which corresponds to the spin concentration obtained for BtuB T188C labeled with M-PROXYL. The outer membranes were diluted to a final concentration of ∼15 µM of BtuB and the same amount of T-CNCbl was added. The samples were transferred to 1.6 mm outer diameter quartz ESR tubes (Suprasil, Wilmad LabGlass) and snap-frozen in liquid nitrogen. Echo-detected field-swept and the phase memory time (*T*_*M*_) experiments were performed using the two pulse Hahn echo sequence *π*/ 2 −*τ*− *π* − *τ*. A Gaussian *π*/ 2 and *π* pulse of 32 ns or 48 ns was used for the Gd(III) or the nitroxide tags respectively, with a 100 ns integration window. For the trityl labels, 64 ns pulses were used, and the echo was integrated for 250 ns. Nitroxide decay curves in a 4-pulse and 5-pulse PELDOR/DEER observer sequences were recorded at 50 K using 48 ns Gaussian pulses. In the 4-pulse decay, the position of the last *π* pulse was increased in steps of 4 ns. In the 5p decay, the position of the two *π* pulses was varied in 4 ns and 12 ns steps, respectively. The *T*_*M*_ and the stretch factor κ were obtained by fitting the normalized Hahn echo decay traces with the stretched exponential function exp[−(2*τ*/ T_M_)^κ^]. For a comparison of the 2-pulse, 4-pulse, and 5-pulse decay traces, the timepoint corresponding to 10 % of initial signal intensity (of the 2-pulse decay) was obtained as mean ± standard deviation (S.D.).

The four-pulse PELDOR/DEER experiments were recorded with a dead-time free sequence *π*/2_*obs*_*− τ*_1_ *− π*_*obs*_ *− t*_1_ *− π*_*pump*_ *−* (*τ* _1_ + *τ*_2_ − *t*_1_) *− π*_*obs*_ *− τ*_2_ using Gaussian pump and observer pulses with a 16-step phase cycling (x[x][x_p_]x) [55, 56]. Nitroxide-nitroxide measurements were acquired at 50 K using 38 ns pump pulse and 48 ns observer pulses. The pump pulse was set to the maximum of the echo-detected field-swept spectrum, while the observer pulses were set at 80 MHz lower and the SRT was kept at 2 ms. For nitroxide-trityl PELDOR, the temperature was set to 50 K when observing the nitroxide and increased to 100 K for observing the trityl. The pump pulse was set to 48 ns or 64 ns when observing the nitroxide at 50 K while the observer pulses were set to 48 ns and a SRT of 2 ms was used. When observing the trityl at 100 K, the pulse lengths were kept at 38 ns and 64 ns for pump and observer pulses, respectively, with a SRT of 3 ms. The frequency offset between the observer and pump pulses was 90 MHz. Nitroxide-Gd(III) PELDOR/DEER experiments were performed at 10 K while pumping the nitroxide and observing the Gd(III). The pump pulse length was 24 ns, while the 48 ns observer pulses were set at 280 MHz lower. The SRT was kept at 1 ms. The deuterium modulations were averaged by increasing the first inter pulse delay by 16 ns for 8 steps for all the samples. The distance distributions were obtained using the MATLAB-based software package DeerAnalysis2018. The form factors F(t)/F(0) obtained after removing the background function from the primary data V(t)/V(0) were fitted to distance distributions by a model-free Tikhonov regularization approach using the L-curve criterion. The suggested regularization parameter (α) was chosen according to the L-curve corner recognition as implemented in the DeerAnalysis software. The error for the probability distribution of distances due to the uncertainty in the background function was determined by a combined variation of the time window and the dimensionality for the spin distribution (see Supplementary Table 2). Additionally, the primary PELDOR/DEER data was analyzed in a combined approach employing deep neural network processing and Tikhonov regularization using the ComparativeDeerAnalyzer program as implemented in the DeerAnalysis2021b software package [57, 58].

The five-pulse PELDOR/DEER experiments were performed according to the pulse sequence *π*/2_*obs*_*−* (/2 − *t*_0_)*− π*_*obs*_ *− t*_0_*− π*_*obs*_ *− t*^′^ − *π*_*pump*_ − (*τ − t*^′^ + *δ*) *− π*_*obs*_ *−* (*τ*_2_ + *δ*). Experiments were performed at 50 K using 48 ns Gaussian observer pulses and a 16-step phase cycling (xx_p_[x][x_p_]x). Nuclear modulation averaging was performed analogous to 4-pulse PELDOR (16 ns shift in 8 steps) with a corresponding shift of the standing pump pulse. The length of the first pump pulse (at ν_obs_ + 80 MHz) was varied (between 30 ns–64 ns as indicated in the corresponding figures) to observe the effect on the amplitude of the 4-pulse artifact. The sech/tanh pulses were designed using EasySpin [59] to sweep over a range of ν_obs_ + 80 MHz ± 33.5 MHz in 400 ns. The truncation parameter β was set to 8/*t*_*p*_. The amplitude and frequency modulation functions of this pulse have been described elsewhere [60, 61]. While using two sech/tanh pump pulses, the moving pulse was frequency swept in a direction opposite to the standing pulse to minimize the excitation time dispersion [22]. Also, this pulse was generated with the four distinct phases for a 16-step phase cycling. The data were analyzed using the Python based DeerLab program [62]. Simulations of rotamer populations and the interspin distances were performed on the calcium and cyanocobalamin bound structure of BtuB (PDB 1NQH) using the rotamer libraries as implemented in the MATLAB-based software package MMM2021.3 [63].

## 3. Results and Discussion

### 3.1 Stability of the spin labels in the cellular environment

Methanethiosulfonate functionalized spin labels give high specificity and reactivity for cysteines. Maleimide reaction with cysteines also exhibits similar specificity at a pH of 6.8–7.0, but overall, the reaction is slower, which could pose some challenge for direct labeling in the extracellular environment. To overcome the limited stability of the common nitroxide labels, their shielded analogues as well as Gd(III) and trityl based labels were introduced [25, 38, 64, 65]. Replacing the *gem*-dimethyl groups with ethyl groups were shown to enhance the stability of the pyrroline nitroxides [38, 66]. The diethyl-shielded MAG1 spin label we used is very similar to the previously reported MAG or M-TEMPO label [32], except that it lacks one double bond, thereby forming a pyrollidine ring instead of a pyrolline ring [54]. MAG was reported to be stable inside *Xenopus laevis* oocytes, in presence of ascorbate, and in the cell lysate of *E. coli* or HEK cells, whereas M-PROXYL and MTSL were shown to be unstable under similar conditions [32, 67, 68]. M-Gd-DOTA and the M-OX063 labels were shown to be stable inside cells [25, 67, 69]. Importantly, none of these labels were used for direct labeling of a membrane protein in *E. coli*. Here we tested M-PROXYL, MAG1, M-Gd-DOTA and the M-OX063 labels (Figure 1A-E) for spin labeling the cobalamin transporter BtuB in both *E. coli* and native OM and compared their performance with the most common nitroxide label MTSL.

**Figure 1.**
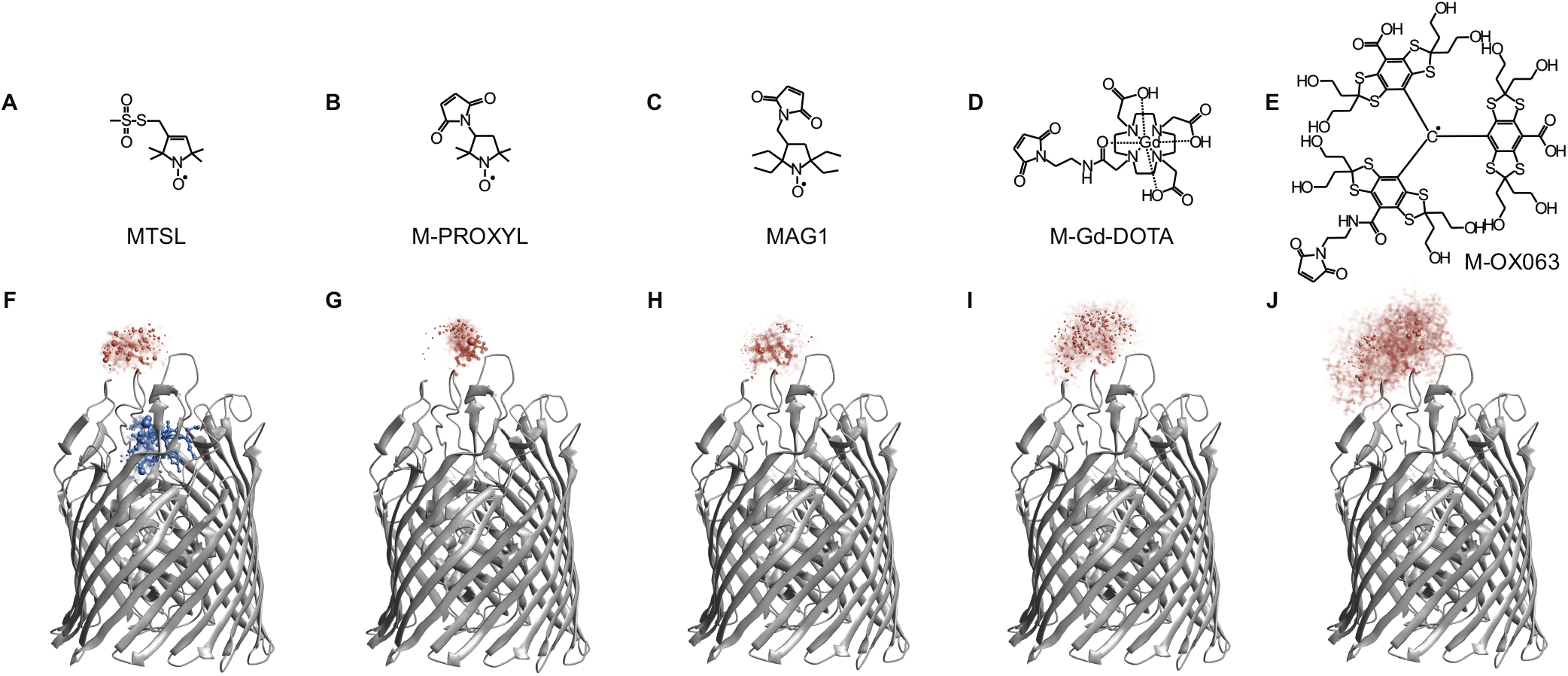
(A-E) The spin labels used in this study and the corresponding rotamer populations at position T188C (F-J), which were simulated on the crystal structure of cyanocobalamin bound BtuB (PDB 1NQH). Rotamer population of TEMPO-attached cyanocobalamin (T-CNCbl, blue, see Figure S5B) is also shown in F. The spheres in the rotameric cloud indicate to the N-O midpoint and their volume corresponds to the relative probability in relation to the total population.

Among these labels, MTSL is the smallest followed by M-PROXYL and MAG1 labels having a rather similar size. The M-Gd-DOTA is larger than the previous ones and the M-OX063 label is the largest. Also, the latter two labels have the same linker, which is much longer when compared with the three nitroxide labels. Simulation of the rotamer populations revealed a cloud of comparable size for MTSL, M-PROXYL, and MAG1. As expected, M-Gd-DOTA occupied a larger cloud and M-OX063 rotamers occupied an even larger conformational space (Figure 1F-J). First, we tested the stability of the labels in presence of ascorbate and in the *E. coli* cell suspension. In line with previous observations [32, 67, 68], both MTSL and M-PROXYL were rapidly reduced in presence of 1–5 mM ascorbate (Figure 2A and S1A) and similar trend was observed in *E. coli* suspension. The slower reduction in the cell suspension (vs. cell lysate [67, 68]) might be due to the lower OD_600_ (0.5) we used as well as that the cell lysate contains more reducing agents. The MAG1 and M-OX063 labels exhibited superior stability in both *E. coli* suspension and 5 mM ascorbate solution (Figure 2). As we reported earlier, OX063 label shows a tendency for aggregation at higher cell density (Figure S1) [50].

**Figure 2.**
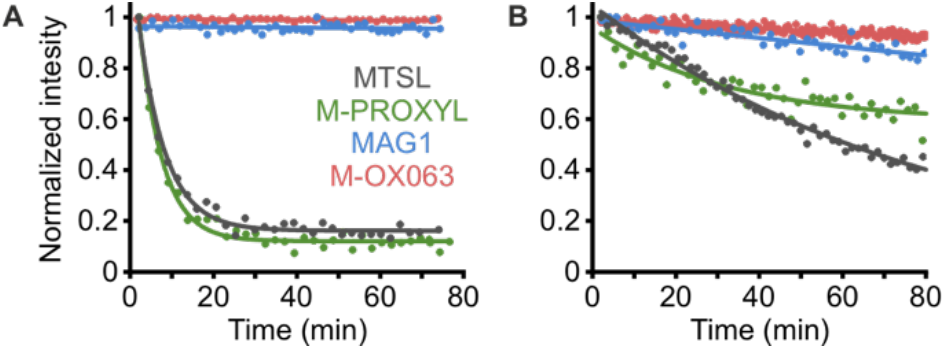
Redox kinetics of the indicated spin labels in ascorbic acid or *E. coli* cell suspension. (A) Reduction of 200 µM spin labels with 5 mM ascorbic acid in MOPS-NaCl (pH 7.5) buffer. (B) Stability of 200 µM spin label in *E. coli* cell suspension in the same buffer at OD_600_ = 5.0 for the nitroxide labels or at OD_600_ = 0.5 for M-OX063 label. The data points were extracted from the double integral of the time resolved spectra and were fitted with either a linear or a mono-exponential function to visualize the overall trend during the observed time window.

### 3.2 Spin labeling in *E. coli* and native membranes

We labeled BtuB at positions T188C and F404C in *E. coli* and native outer membranes using these labels (Figures 1, 3, and S2). Expression analysis using SDS-PAGE revealed clear bands corresponding to BtuB in *E. coli* cells (in both RK5016 and BL21 strains) and native membranes (Figure S3). To maintain cell viability, we limited the labeling time to 1 h at RT. Despite the short incubation time, M-PROXYL and MAG1 gave signals comparable with MTSL in both the environments (Supplementary Table 1). Yet, for more restricted sites (see V90C in Figure S2 and S4), MTSL would be a better choice owing to its smaller size. The higher stability of the MAG1 in the cell suspension (Figure 1B) did not lead to an increase in signal in *E. coli*. This observation suggests that the spin label reduction *per se* may not have a direct effect on labeling on the cell surface. The exact location for the reduction remains unknown. Previously, we showed that the supernatant from the cell suspension has no effect, whereas heat inactivation of the cells considerably decreased MTSL reduction, suggesting the involvement of an active process [50]. The M-OX063 label also gave signals in both the environments, but lower than the nitroxides. Overall, the amount of labeling is comparable with the MTS-OX063 which we reported previously (Figure S5A) [50]. Most importantly, the background labeling was significantly reduced for M-OX063. Similar results were obtained upon labeling the position F404C using these labels (Figure S6 and Supplementary Table 1). In general, the background labeling is increased in the outer membranes, possibly due to an enhanced interaction between the labels and the membrane surface. Labeling using M-Gd-DOTA did not work in *E. coli* and further experiments suggested that Gd(III) might be released from the M-DOTA (Figure S7). It has been shown that an azide functionalized Gd-DOTA can label eGFP inside *E. coli* when used at a higher (1 mM) concentration [70]. It appears that at the low concentration we used (15 µM), the stability of the label might be compromised.

**Figure 3.**
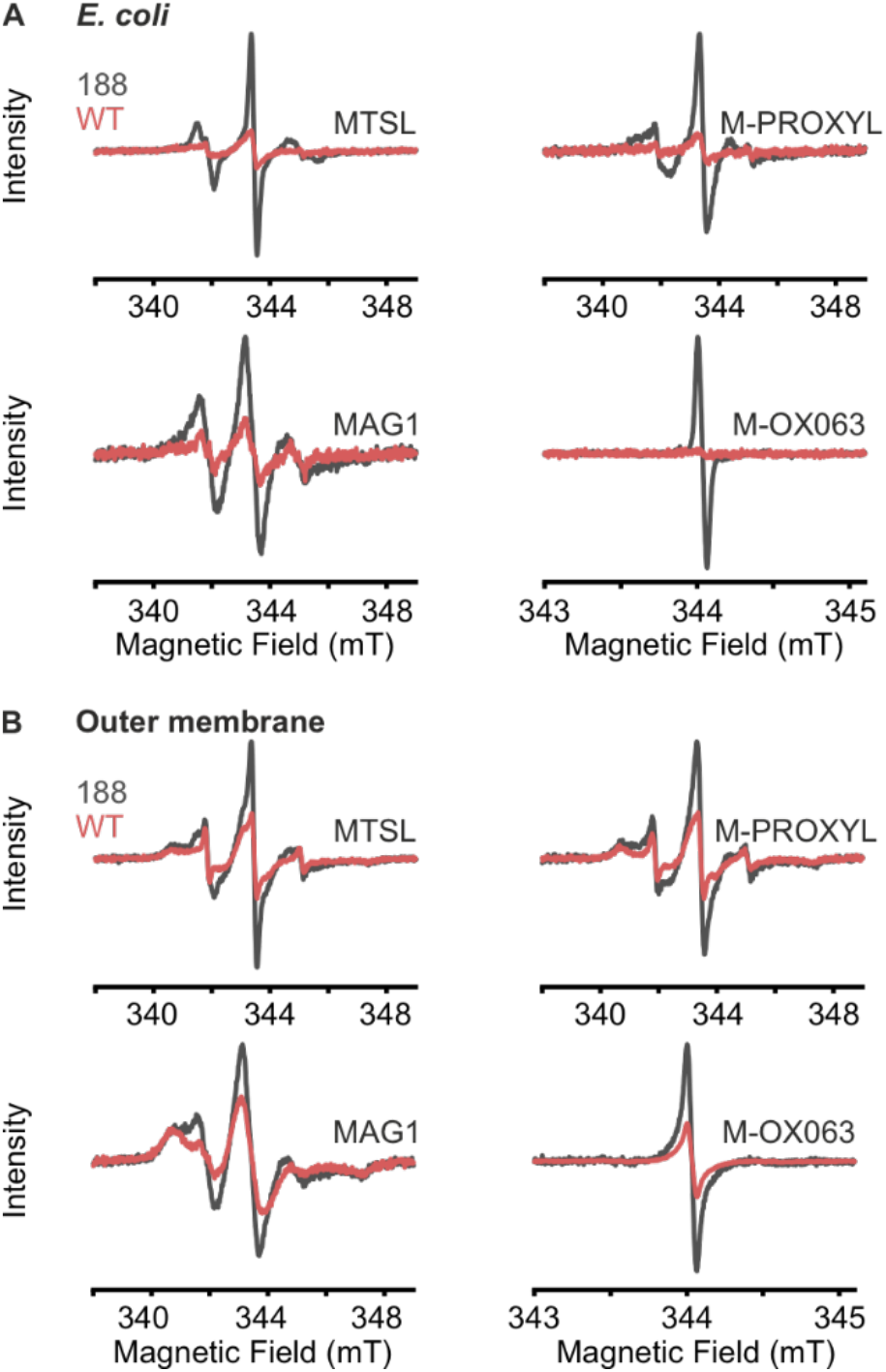
*In situ* room temperature continuous wave (cw) ESR spectra for nitroxide and M-OX063 spin labels. (A) Spin labeling of BtuB wild-type (red) or T188C (grey) in whole *E. coli* cells, or (B) in isolated native outer membranes. The spin concentrations are given in Supplementary Table 1.

### 3.3 Transversal relaxation in the native environments

We determined the phase memory time (*T*_*M*_) for all the labels after attaching to BtuB in *E. coli* and OM (Figure 4). In *E. coli* at 50 K, M-PROXYL and M-OX063 gave the longest *T*_*M*_ of ∼4.0 µs (Table 1). MTSL and MAG1 gave a *T*_*M*_ of ∼3.0 µs. These values are in the range typically observed for solvent-exposed sites [15]. In such a case, the curves would decay according to the function exp[−(2*τ*/ *T*_*M*_)^κ^]. Fitting the curves with this function gave values for κ = 1.0 (mono-exponential decay) for MTSL and MAG1, and 1.3 and 1.5 for M-PROXYL and M-OX063, respectively. Moving to 100 K, the *T*_*M*_ was reduced by more than two-fold with a further decrease of the κ values for all the nitroxides. Such small values of κ reveal the involvement of a dominant dynamical process (in addition to the solvent protons), which has a rate comparable with the spin echo decay. Rotating methyl groups are well documented to account for such a process, with a range of *T*_*M*_ (0.4–5.0 µs) and κ (0.8–2.2) values depending on the type of the methyl protons in the environment [15]. The second loop carrying the spin label would be integrated into the core oligosaccharides of the LPS molecules, which forms the outer leaflet of the membrane. It is possible that this unique environment makes a major contri-bution to the relaxation behavior observed here. For the M-OX063 label, the average distance of closest approach for protons to the central carbon might be considerably increased, resulting in a temperature independent *T*_*M*_ in the range of 50−100 K. Interestingly, in the outer membrane, the *T*_*M*_ was significantly reduced for all the labels to the range of 2.0–2.5 µs at 50 K. This would not be expected, and a fully quantitative interpretation of this observation is not feasible as the membranes contain higher background labeling. These labels adsorb or even aggregate in the membrane, and they might have a shorter *T*_*M*_ as compared to the labels attached with the protein [49]. As in *E. coli* suspension, moving to 100 K reduced the *T*_*M*_ by more than two-fold accompanied with a further reduction of κ for the nitroxides, whereas the M-OX063 label showed little effect. Owing to the faster relaxation, M-Gd-DOTA was measured at 10 K (and only in the OM as labeling could not be achieved in *E. coli* suspension), which gave a *T*_*M*_ of ∼2.7 µs with κ = 1.0. In summary, M-PROXYL gave the longest *T*_*M*_ among the nitroxides, whereas the M-OX063 label has the longest *T*_*M*_ among all the labels, which is independent of the temperature between 50 and 100 K.

**Table 1.** Phase memory time (*T*_*M*_) and stretch factor κ determined from fitting the 2-pulse decay curves in *E. coli* cells and native outer membranes using the function V(2*τ*) = A(0)·exp[−(2*τ*/ T_M_)^κ^]. Spin labels were attached to position T188C in BtuB and measurements were performed at 10 K, 50 K, or 100 K as indicated. The experimental data and the corresponding fits are shown in Figure S8. The error values for *T*_*M*_ and κ were below 1 % and are therefore not shown.

| Spin label | $T_M$ Outer membrane (μs) | | | $\kappa$ Outer membrane | | | $T_M$ *E. coli* (μs) | | $\kappa$ *E. coli* | |
| --- | --- | --- | --- | --- | --- | --- | --- | --- | --- | --- |
|  | 10 K | 50 K | 100 K | 10 K | 50 K | 100 K | 50 K | 100 K | 50 K | 100 K |
| MTSL | - | 2.0 | 0.8 | - | 0.8 | 0.7 | 3.0 | 1.2 | 1.0 | 0.8 |
| M-PROXYL | - | 2.4 | 1.1 | - | 0.9 | 0.8 | 4.1 | 1.6 | 1.3 | 0.8 |
| MAG1 | - | 2.3 | 0.7 | - | 0.8 | 0.7 | 3.0 | 0.8 | 1.0 | 0.7 |
| M-OX063 | - | 2.5 | 2.5 | - | 1.0 | 1.0 | 4.2 | 4.1 | 1.5 | 1.4 |
| M-Gd-DOTA | 2.6 | - | - | 1.0 | - | - | - | - | - | - |

**Figure 4.**
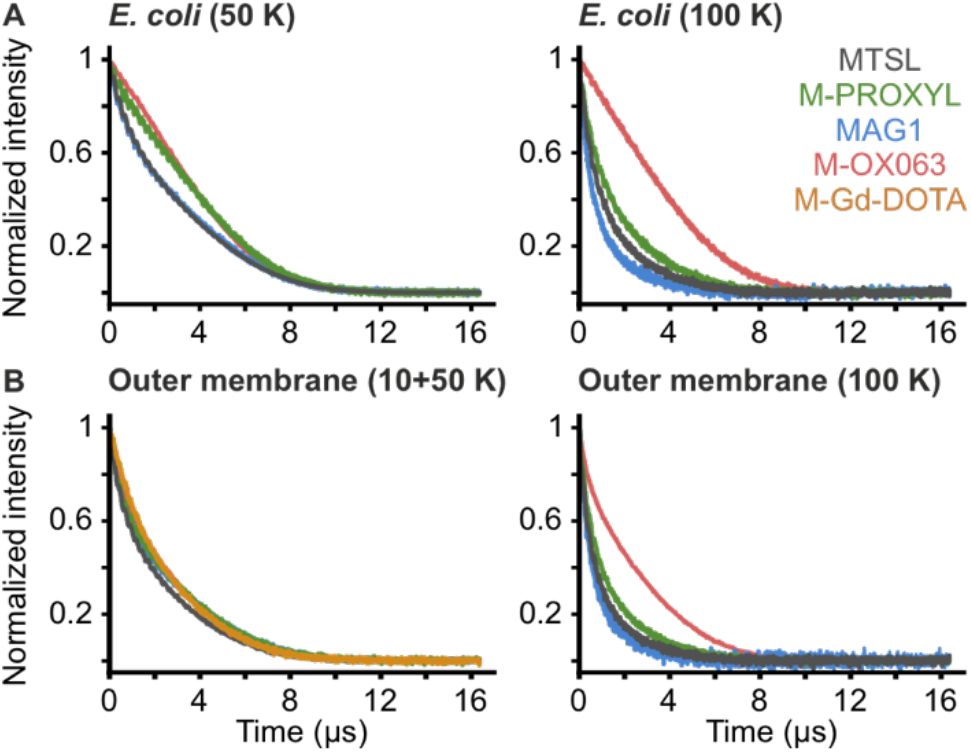
*In situ* analysis of transversal relaxation for nitroxides, trityl, and Gd(III)-based spin labels attached to BtuB T188C. (A) Normalized Hahn echo decay curves for nitroxides and trityl at 50 K (left) or 100 K (right) are shown. (B) Relaxation curves for the nitroxides and trityl labels at 50 K and for M-Gd-DOTA at 10 K (left) or for nitroxides and trityl at 100 K (right) in isolated native outer membranes. Further analysis of the data is shown in Figure S8 and values for *T*_*M*_ and the stretch factor *k* are given in Table 1.

### 3.4 Distance measurements using 4-pulse PELDOR

In the echo-detected field-sweep (FS) spectra at the Q-band frequency, the maxima of the nitroxide and the M-OX063 labels are separated by ∼90 MHz. This separation increases to ∼280 MHz for M-Gd-DOTA (Figure 5). For the PELDOR experiments, we attached the labels at position T188C on the second extracellular loop in BtuB and determined the distances to the spin labeled substrate (T-CNCbl, Figure S5B). For Gd(III)-NO PELDOR, the pump pulse was placed on the nitroxide. For the M-OX063-nitroxide pair, we pumped either of them independently. As we reported earlier, the singly labeled variant T188C revealed a stretched exponential decay devoid of any distances (Figure S9). Despite the large size difference between the labels, simulations predicted distance distributions with a comparable width. The MTSL gave the narrowest distribution (2.51±0.25 nm) with a major peak and a shoulder, and significantly narrower than the simulation (Figure 6A). M-PROXYL and MAG1 gave distance distributions with an overall width slightly broader than MTSL (2.37±0.44 nm and 2.41±0.40 nm, respectively), with a more pronounced contribution of the shoulder peak (Figure 6B-C). Yet, as we mentioned earlier, for a restricted site MTSL gives better performance, at which even M-PROXYL failed to produce any appreciable labeling (Figure S10). The RK5016 *E. coli* strain routinely used for BtuB expression lacks the O-antigen oligosaccharides, which are attached to the lipid A core of the LPS. Measurements using M-PROXYL in the commercial BL21(DE3) strain, which carries the O-antigen gave identical distances, revealing that the O-antigen *per se* does not have an influence on the conformation of the second extracellular loop (Figure S12).

**Figure 5.**
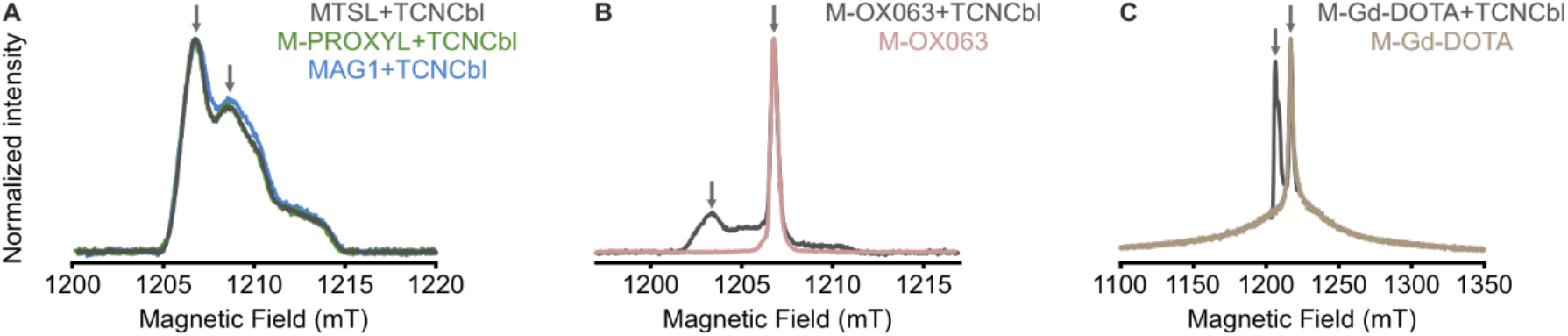
Echo-detected field-swept spectra of spin labels attached to BtuB at T188C in isolated native outer membranes. (A) Normalized spectra of nitroxide spin labels at 50 K, (B) of M-OX063 at 50 K, and (C) of M-Gd-DOTA at 10 K. Spectra were acquired in the presence or absence of the spin labeled substrate (T-CNCbl). Arrows indicate the frequency offset of 80 MHz for nitroxide-nitroxide, 90 MHz for nitroxide-trityl, and 280 MHz for nitroxide-Gd(III) PELDOR/DEER measurements.

**Figure 6.**
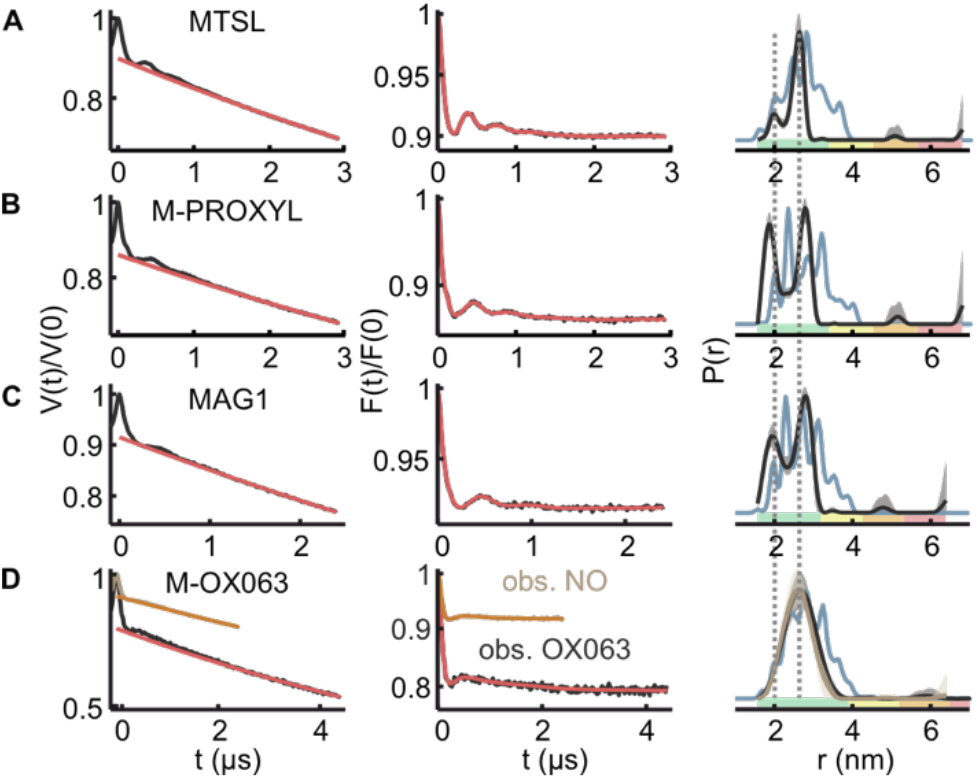
PELDOR data for nitroxides and M-OX063 labels attached to BtuB T188C in whole *E. coli* cells. Distances were measured to the bound spin labeled substrate (T-CNCbl). (A-D) Left panels show the primary data (in grey or light brown) with the intermolecular (background) contribution overlaid (red or orange); Middle panels represent the background corrected form factors overlaid with the corresponding fit; Right panels show the calculated distance distribution using Tikhonov regularization. The error bars represent variations of the probability distribution corresponding to the uncertainty in the background function (see Supplementary Table 2). The reliability of shape, width, and mean distance are represented with the color codes as implemented in the DeerAnalysis software. Measurements were performed at 50 K or 100 K for the nitroxides or the M-OX063, respectively. The corresponding simulations of the probability distributions (PDB 1NQH) are overlaid in light blue. The L-curves are given in Figure S11.

The C-S bond of the maleimide labels forms a chiral center, which could lead to the orientation of the spin label as two epimers [68]. Thus, the shoulder may also arise from a distinct subset of rotamers. Interestingly, this bimodal distribution was absent for the M-OX063 label (Figure 6D). The overall width of the distribution (2.54±0.29 nm) is rather similar to the nitroxide labels, implying that the electron density is mostly localized on the central carbon atom. To further elucidate these observations, we labeled position F404C with MTSL, M-PROXYL, and M-OX063 and determined the distances to T-CNCbl (Figure S13-14). These experiments revealed a single distance peak (at 2.04±0.27 nm, 2.09±0.32 nm, and 2.18±0.32 nm, respectively) with a very similar mean and width. Thus, the bimodal distribution for M-PROXYL and MAG1 labels at position T188C might be mediated through the interaction with surroundings (either two distinct set of rotamers or loop conformation). Overall, all the distance distributions were least affected when determined in the isolated native outer membranes (Figure 7). For M-OX063 in *E. coli*, observing the NO (T-CNCbl) gave a smaller modulation depth (∼9% vs.∼20% while observing M-OX063, Figure 6D), in agreement with an overall lower labeling efficiency when compared to the other labels. In the membranes as well, observing NO gave a similar value (∼9%), whereas observing M-OX063 did not increase it further, likely due to the higher background labeling (Figure 7D).

**Figure 7.**
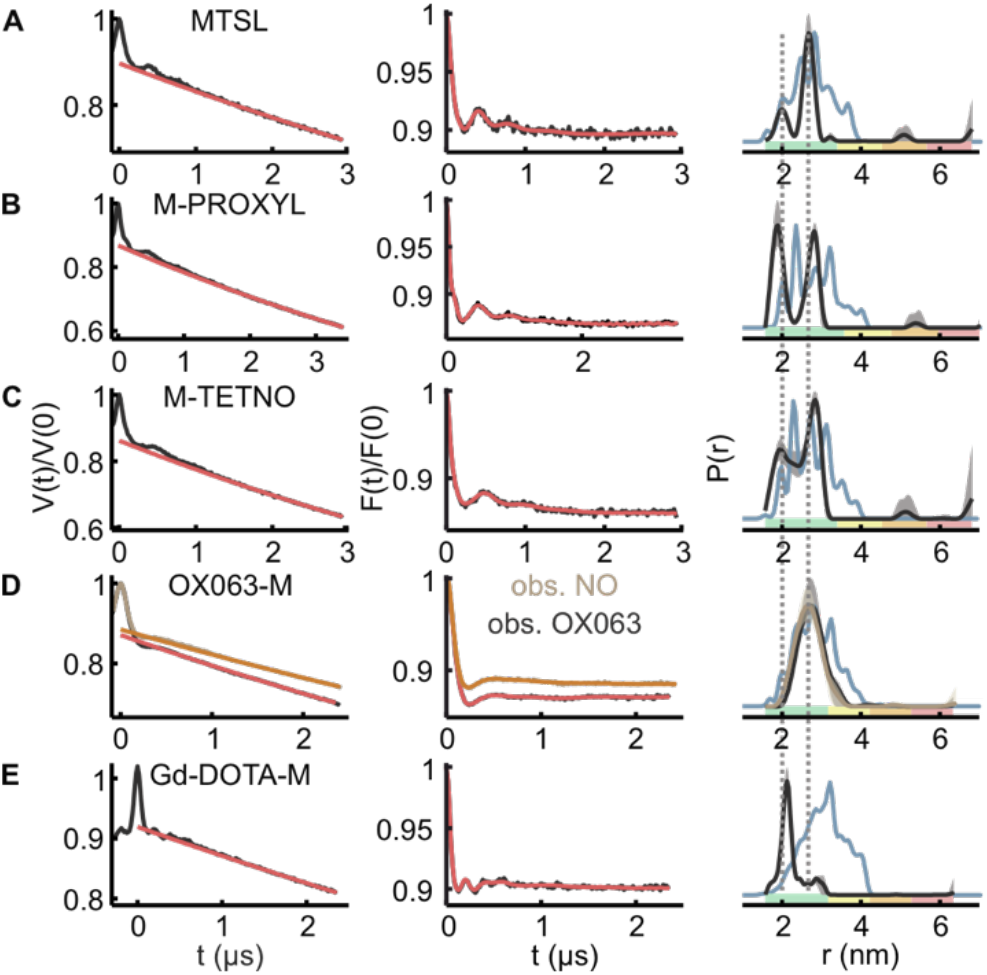
PELDOR data for nitroxide, M-OX063, and M-Gd-DOTA labels attached to BtuB T188C in the native outer membranes. Distances were measured to the bound spin labeled substrate (T-CNCbl). (A-D) Panel description as in Figure 6. (E) Data for M-Gd-DOTA obtained at 10 K.

Owing to the instability of M-Gd-DOTA in the *E. coli* suspension (Figure S7), we focused on the OM preparation for the PELDOR experiments. The simulation predicted a broad distribution of the rotamers (Figure 1I) and the interspin distances. Surprisingly, the PELDOR experiments revealed a narrow distribution centered at 2.26±0.36 nm (Figure 7E). Thus, the native membrane environment considerably altered the rotamer distribution leading to the selection of a particular set of rotamers. As the linker is the same as for the M-OX063 label, it is likely that the interaction of the central tetraaza ring of the DOTA with the surroundings plays an important role for the narrowing of the distance distribution. Earlier we reported a similar observation where the MTS-TAM1 showed a much narrower distribution as compared to the MTS-OX063 label, despite having a very similar linker [50]. The overall modulation depth is ∼10%, which is similar to the value obtained for the other labels. We did not perform a quantitative analysis of the sensitivity for the different orthogonal pairs. Nevertheless, as we reported earlier [50], observing OX063 at 100 K might give a clear advantage over other labels as the narrow line gives a much larger signal and the experiment can be repeated faster (SRT of ∼2 ms). An independent analysis of the data sets using deep neural network analysis and the ComparativeDeerAnalyzer gave similar distance distributions (Figures S15-16).

### 3.5 5-pulse PELDOR on membrane proteins in detergent micelles, native membranes, and *E. coli*

The dynamical decoupling of electron spins by Carr-Purcell pulse sequence was shown to prolong the coherence lifetime even in the lipid environments [24]. Here, the spin labeled WALP23 peptide reconstituted into a DOPC bilayer revealed a mono-exponential decay of the phase coherence, yet the 5-pulse PELDOR observer sequence increased the time at which the initial echo amplitude has decayed to 10% by a factor of ∼1.9. Whether such a gain can be obtained for a spin labeled membrane protein in the native environments is yet to be elucidated. Here, we first tested the performance of shaped inversion pulses for the suppression of the partial excitation artefact using the detergent (DDM) solubilized ATP-binding cassette transporter LptB_2_FG [71]. It transports lipopolysaccharide from the inner membrane of Gram-negative bacteria at the expense of ATP binding and hydrolysis [72]. We spin labeled position V18C located in the nucleotide binding domains (NBDs), which are solvent accessible (Figure 8A-B). The 2-pulse Hahn echo sequence and the 4-pulse observer sequence revealed that the echo decays to 10% of the intensity at ∼6.4 µs (in buffer containing 15% d_8_-glycerol). Interestingly, the symmetrized Carr-Purcell 5-pulse sequences increased this value to 12.9 µs and by a factor of ∼1.9. Similar enhancement has been achieved for the detergent solubilized betaine transporter BetP using a 7-pulse PELDOR observer sequence [73].

**Figure 8.**
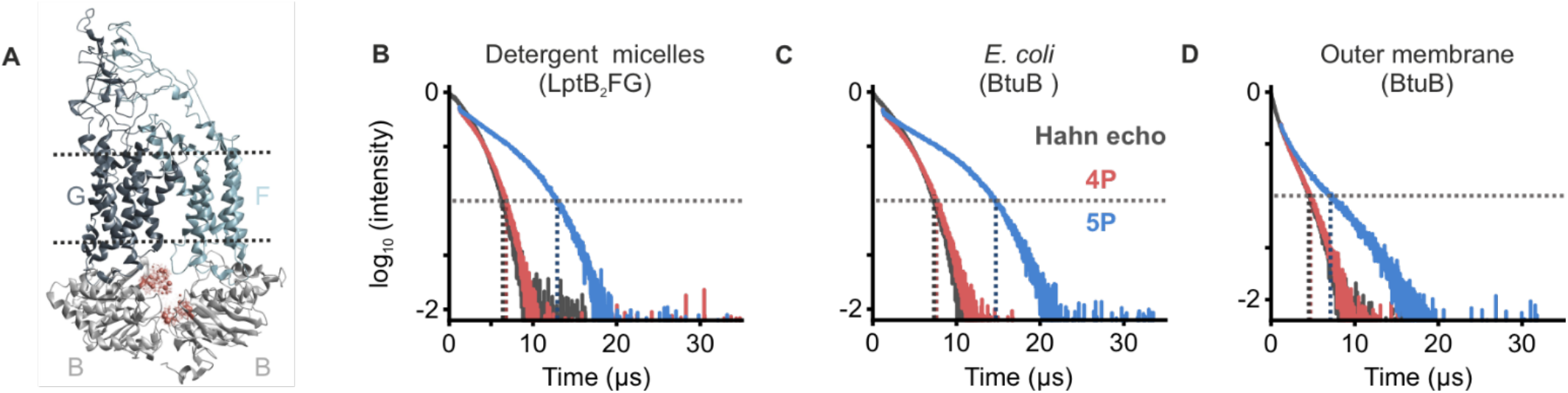
Decay traces of a Hahn echo, four-pulse, and five-pulse PELDOR/DEER observer sequence in DDM micelles, *E. coli* cells, or native outer membranes at 50 K. (A) Cryo-EM structure of LptB_2_FG (PDB 6MHU) with the rotamers for MTSL (in red) at position LptB-V18C. (B) 2-pulse (grey), 4-pulse (red), and 5-pulse (blue) decay traces obtained for MTSL labeled LptB-V18C in DDM micelles. Dashed lines indicate the 10 % of the Hahn echo amplitude, which gave 6.4 µs for the 2-pulse and 6.8 µs for the 4-pulse observer sequences. The 5-pulse sequence prolongs the decay by a factor of ∼1.9 (to 12.9 µs) as compared to the 4-pulse observer sequence. (C) Decay traces for BtuB T188C variant labeled with M-PROXYL in the presence of spin labeled substrate (T-CNCbl) in whole *E. coli* cells, which gave values of 7.3 µs, 7.6 µs, and 14.7 µs for the 2-pulse (grey), 4-pulse (red), and 5-pulse (blue) sequence, respectively, prolonging the echo decay by a factor of ∼1.9 for the 5-pulse sequence. (D) Echo decay curves measured on MTSL labeled BtuB T426C with bound T-CNCbl in native outer membranes. The 10 % intensity values are 4.5 µs, 4.7 µs, and 7.1 µs for the 2-pulse, 4-pulse, and 5-pulse observer sequence, respectively, which corresponds to an increase of ∼1.5 for the 5-pulse sequence.

Initially, we tested the performance of Gaussian pulses having different amplitudes for the standing and moving pump pulses on artifact suppression (Figure 9B-D). Applying a standing pump pulse with a broader excitation bandwidth was shown to decrease the amplitude of the artifact [17, 22, 24]. When we applied a 38 ns standing and a 48 ns moving Gaussian pulse, the data contained a large partial excitation artifact. When the standing pump pulse was set to 30 ns, the amplitude of the artifact (with respect to the principal dipolar signal) was reduced by a factor of ∼2 (as calculated using the DeerLab program). Applying a 400 ns symmetric sech/tanh standing pump pulse (at ν_*pump*_ = ν_*obs*_ + 80 MHz ± 33.5 MHz) reduced the artifact by a factor of ∼7. Though the sech/tanh pulse we used is rather long, in general, these pulses invert the spin packets in time window much shorter than the pulse length [21]. Thus, least interference is expected if the pulse length is smaller than the period of the oscillation corresponding to the perpendicular orientation of the interspin vector (θ = 90). Comparison of the dipolar spectra between 4-pulse and 5-pulse experiments revealed good agreement (Figure 9G, left), revealing that the pulse has none or only modest effect on the distances in the range determined (≥ 3 nm).

**Figure 9.**
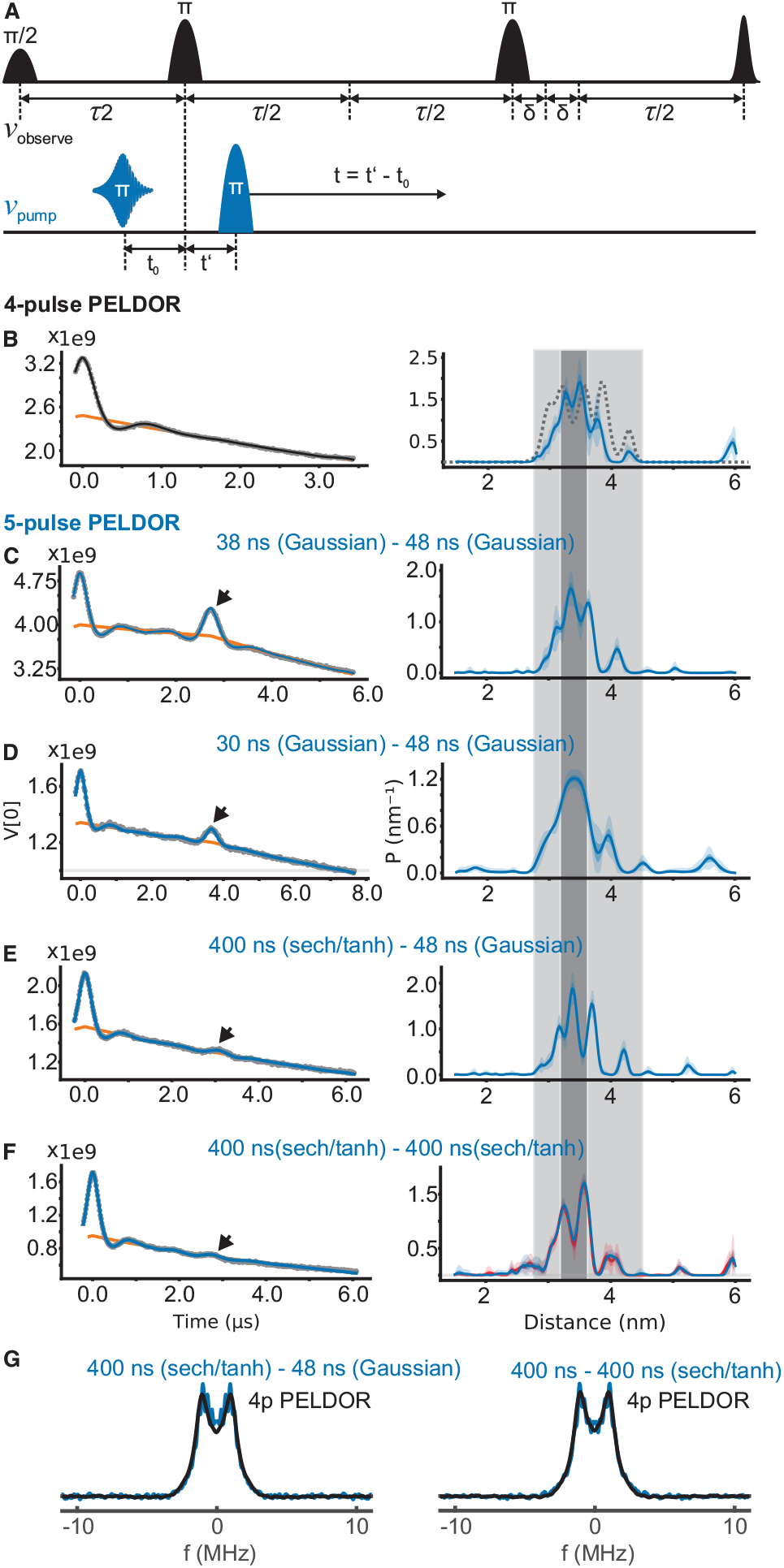
Five-pulse PELDOR measurements on the detergent solubilized lipopolysaccharide transporter LptB_2_FG. (A) The forward five-pulse PELDOR sequence. (B) The 4-pulse PELDOR data (left) and the corresponding distance distribution (right). The range corresponding to the *r*_*max*_ is highlighted in dark grey and distances spanning the simulation (PDB 6MHU) are highlighted in light grey for a direct comparison with the output from the 5-pulse data. (C-F) 5-pulse PELDOR data (in grey) obtained with different standing and moving pump pulses as indicated. The 4-pulse artifact is indicated with an arrow and it is time shifted between different data corresponding to the observed dipolar evolution time. The data were fitted (in blue) by including the 4-pulse artifact using the DeerLab program. The background function is overlaid (in orange) and the resulting distribution is shown on the right. For the data presented in (F), an additional analysis was performed as a 4-pulse PELDOR data (in red), which gave identical distribution. (G) The dipolar spectrum obtained from 4-pulse PELDOR is overlaid with the corresponding data from the 5-pulse PELDOR experiment as indicated.

We further tested the performance of two successive sech/tanh pump pulses (Figure 9F). Here we frequency swept the moving pump pulse in the opposite direction to compensate for the dispersion of the excitation time introduced by the first pump pulse [22]. This setup further suppressed the artifact. Such a suppression could not be achieved in a previous study employing asymmetric sech/tanh pulses, which can provide offset-independent adiabaticity where the resonator bandwidth is limiting a broad excitation [24]. The differences in the pulse parameters (length, bandwidth etc.) and the experimental settings (amplifier and the resonator) might account for the increased artifact suppression we achieved. For an overall better sensitivity, we placed the pump pulses at +40 MHz from the center of a fully over coupled resonator mode, where we observed an inversion (*M*_*z*_*/M*_*0*_) of ∼-0.73. This corresponds to an inversion efficiency *I* = [1 − *M*_z_/*M*_0_]/2 of ∼0.87. The small artifact level suggests that this inversion efficiency might be achieved over the relatively small bandwidth of the pulse (± 33.5 MHz). This setup also gave a modulation depth, which is ∼2x larger. The analysis fitted a zero amplitude for the artifact, suggesting that at this level the artifact has minimum effect on the output. The dipolar spectrum perfectly overlays with the 4p-PELDOR spectrum (Figure 9G, right). Thus, the small differences in the distance distribution might be due to the differences in the noise level. Overall, the 5-pulse PELDOR data independent of the amplitude of the artifact gave a probability distribution similar to the 4-pulse PELDOR and the distances span the range predicted by the simulation.

### 3.6 Trityl-nitroxide 5-pulse PELDOR in the native membrane environment

Most of the 5-pulse experiments so far have been performed with the NO-NO system. Spindler *et al*. reported the experiment on a Co(II)-NO system and showed that employing two sech/tanh pump pulses can considerably suppress the 4-pulse artifact [22]. Even though a standing pump pulse with a broader excitation bandwidth was employed in their study, a complete suppression of the artifact could not be achieved. Trityl labels offer a unique advantage with their narrow linewidth. It can be totally excited with a Gaussian pulse, which, in the linear regime also has a Gaussian excitation profile. Before attempting such an experiment, first we characterized the dynamical decoupling in the cellular environment using BtuB T188C variant labeled with M-PROXYL. With the 5-pulse PELDOR observer sequence, the echo decayed much slower so that the time to reach the 10% of the 2-pulse echo intensity was prolonged by a factor of ∼1.9 (from 7.6 µs to 14.7 µs, Figure 8C). Similar experiment in the isolated membranes using MTSL (BtuB-T426R1+T-CNCbl) revealed an enhancement by a factor of ∼1.5 (Figure 8D). A somewhat reduced enhancement might in part be attributed to the background signals, which adsorb with the membrane and therefore would have a reduced *T*_*M*_.

Next, we performed 5-pulse PELDOR experiment on BtuB labeled with the OX063 tag on its periplasmic turn at position T426C (Figure S2) in presence of T-CNCbl in the isolated native outer membranes. Here we used a MTS functionalized OX063 (for the reason of limited availability of M-OX063), which we had previously reported (Figure S5A) [50]. We observed nitroxide (T-CNCbl), while pumping the narrow trityl line with Gaussian pulses. A 4-pulse PELDOR experiment on this sample enabled the observation of dipolar evolution for 4 µs in an overnight measurement (∼12 h, Figure 10A). It is insufficient to observe the cross-membrane distances and correspondingly, the distribution revealed large error bounds. We performed a 5-pulse PELDOR experiment on this sample using 38 ns (standing) and 64 ns (moving) Gaussian pulses. A 6 µs long dipolar evolution could be observed within the same time, which enabled the observation of the distances with higher confidence (Figure 10B). Remarkably, this setup was sufficient to totally suppress the partial excitation artifact and the data perfectly overlaid with the output from the 4-pulse PELDOR experiment. This also reveals that the 16-step phase cycling effectively suppresses unwanted spin echoes generated by the coherent pulses [56]. Another experiment employing two 64 ns pump pulses increased the amplitude of the artifact similar to the total signal (data not shown). To characterize the size of artifact in more detail under these pulse settings, we performed a similar experiment after attaching the M-OX063 label at position T188C. The spin pair has a mean distance distribution of 2.18±0.32 nm (Figure 6B), which separated the artifact well beyond the dipolar oscillations (Figure 10C). As evident, the artifact is suppressed to typical level of noise in PELDOR experiments, such that the distances can be extracted using existing tools for 4-pulse PELDOR data analysis.

**Figure 10.**
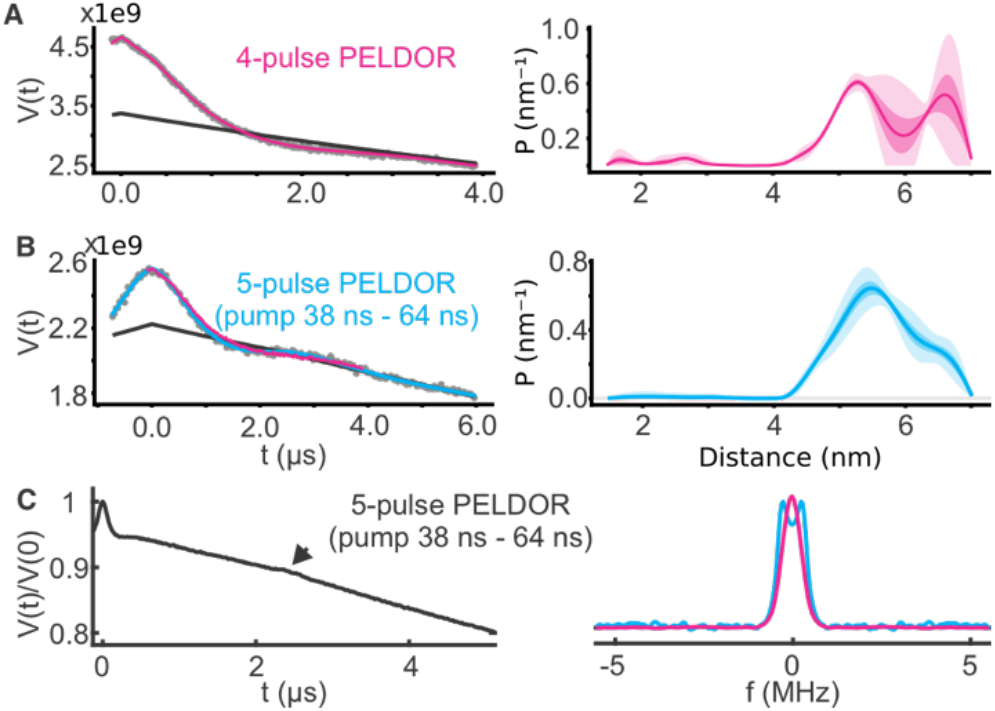
Five-pulse PELDOR/DEER measurements for BtuB T426C and T188C labeled with MTS-OX063 and M-OX063, respectively. Distances were determined to the bound T-CNCbl using all Gaussian pulses and pumping OX063. (A) The primary data from 4-pulse (in grey) and 5-pulse PELDOR (B, in grey) with the corresponding distances on the right. The corresponding fits obtained using the DeerLab program is overlaid in magenta or cyan. (C) The 5-pulse PELDOR experiment between BtuB T188C labeled with M-OX063 and the bound T-CNCbl (left) and the dipolar spectra corresponding to (A) and (B) are overlaid in the right panel.

## 4. Conclusions and Outlook

Under the reducing cellular environments, nitroxides and MTS functionalized labels are unstable. To overcome these problems, maleimide functionalized (shielded) nitroxide-, Gd(III)-, and trityl labels were employed [32, 50, 64, 67-69]. In this study, we explored the feasibility of using maleimide functionalized Gd(III), nitroxide, and trityl spin labels for direct labeling of the cobalamin transporter BtuB in *E. coli* and isolated native membranes. We performed distance measurements between these labels to the spin labeled substrate, T-CNCbl. Overall, the performance of these nitroxide and trityl labels is comparable with their MTS functionalized analogues we reported earlier [44, 50] and the application of Gd(III) in the outer membrane environment is demonstrated for the first time.

In part, these studies were motivated by our previous observation that *E. coli* actively reduced MTSL in a cell density dependent manner over time [50]. It is unclear where exactly the reduction occurs (extracellular space, outer membrane, periplasm, inner membrane, or the cytoplasm), but it appears that this process as such might not have a significant effect on the labeling itself. A fast transversal spin relaxation is the major limitation for measuring longer distances in membrane proteins, especially in the membrane environment. We show that the Carr-Purcell 5-pulse PELDOR can significantly prolong the observable dipolar evolution for membrane proteins, even in the cellular and membrane environment. This offers the possibility to observe the dipolar evolution time up to 12.5 µs in *E. coli* without protein or lipid deuteriation (Figure 8C). Also, the partial excitation artifact in a NO-NO measurement is substantially suppressed while using sech/tanh pump pulses. Further, in a trityl-NO case, simple Gaussian pump pulses of varying amplitude are sufficient to suppress the artifact to the typical noise level. Considering the different tools available for the analysis of the 5-pulse data, a total suppression of the artifact would not be necessary [62, 74]. Yet, the sech/tanh pulses provide better artifact suppression and higher modulation depth, albeit costing a reduction in the observed dipolar evolution corresponding to the length of the moving pump pulse (∼ ≤ 400 ns).

Spectroscopically orthogonal labels can provide more information per sample as compared to an identical spin pair. Following a first demonstration of this possibility using ^14^N/^15^N labeling, several examples including the triple labeling using Gd(III), Mn(II), and NO tags have been reported [75-83]. In the Gd(III)-NO case, faster relaxation and a higher transition moment for Gd(III) enable the spectroscopic separation. For the NO-trityl case, the orthogonality comes from the resonance separation of the narrow intense trityl line towards the edge of the nitroxide spectrum. Such a combination of different labels is a potential tool to study heterooligomeric complexes. At the most commonly used Q-band frequency, the Gd(III)–NO– trityl tags make an ideal orthogonal three spin system. Our results open the possibility to perform such experiments on complex membrane proteins in the native environments.

## Declaration of competing financial interests

The authors declare no competing financial interest.

## Acknowledgements

This work was financially supported through the Emmy Noether program (JO 1428/1-1) and a large equipment funding (438280639) from the Deutsche Forschungsgemeinschaft and the Science Funding from Johanna Quandt Young Academy at Goethe to B.J. Synthesis of OX063 labels was supported by Russian Ministry of Science and Higher Education grant to V.M.T and E.G.B. (14.W03.31.0034). We thank Alberto Collauto and Luis Fábregas Ibáñez for fruitful discussions, Sunil Saxena and Zikri Hasanbasri for providing the PDB file of M-OX063 optimized for MtsslWizard, and Gunnar Jeschke for creating the rotamer library of M-OX063 and a critical reading of the manuscript.

## Supplementary material

Supplementary information associated with this article is provided.

## Notes

### Competing Interest Statement

The authors have declared no competing interest.

