## Supplemental Information for "*In situ* distance measurements in a membrane transporter using maleimide functionalized orthogonal spin labels and 5-pulse electron double resonance spectroscopy"

### Supplementary Figures

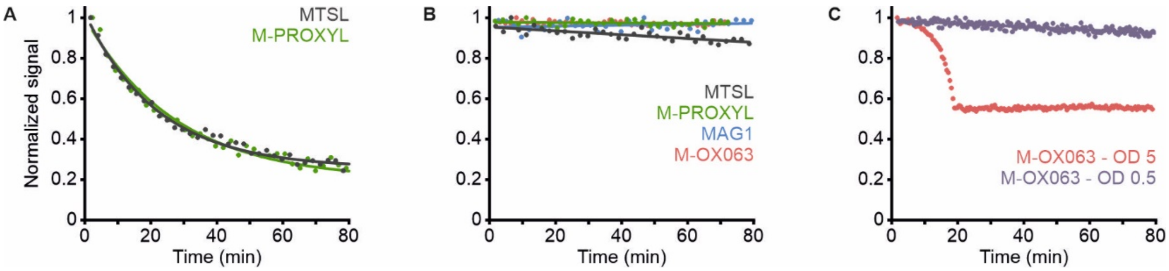

**Figure S1.** Decay curves of indicated spin labels in buffer, ascorbic acid, and *E. coli* cell suspension. (A) Reduction of 200 µM spin label with 1 mM ascorbic acid in MOPS-NaCl (pH 7.5) buffer. (B) Stability of 200 µM spin label in MOPS-NaCl buffer (pH 7.5), and (C) of 200 µM M-OX063 in *E. coli* cell suspension at $OD_{600}$ = 0.5 (purple) or 5.0 (red). At higher cell densities, M-OX063 shows a tendency for aggregation as evident from a steep decline of the signal intensity during the early part of the curve. The double integral is shown as a function of time with a linear or mono-exponential fit to visualize overall trend.

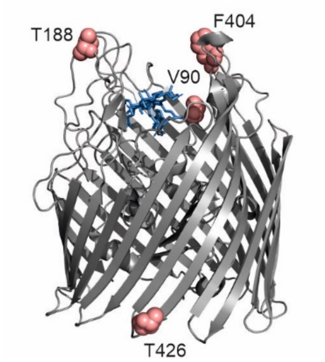

**Figure S2.** Crystal structure of BtuB (PDB 1NQH) with the spin labeled positions and the bound cyanocobalamin substrate highlighted (CNCbl, in blue). Positions T188 and F404 are located on extracellular loops 2 and 7, respectively, V90 on the plug domain of the protein, and T426 is located on a periplasmic turn.

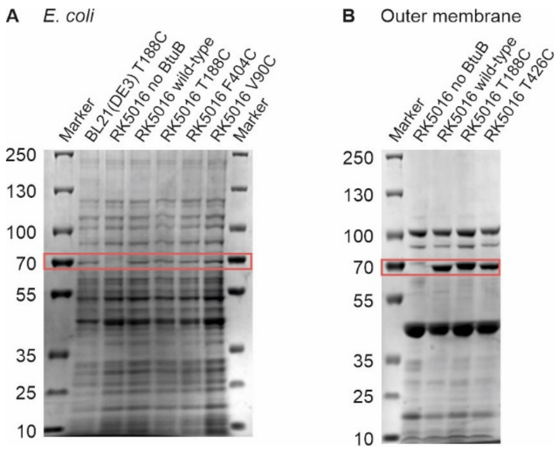

**Figure S3.** Characterization of BtuB variants in *E. coli* and native outer membranes. (A) SDS-PAGE showing the expression of BtuB wild-type and the cysteine variants (∼ 68 kDa, highlighted inside the red box) in RK5016 and BL21(DE3) *E. coli* cells. The band corresponding to BtuB was confirmed by growing RK5016 cells in the absence of the plasmid (indicated as RK5016 no BtuB). (B) SDS-PAGE showing the BtuB wild-type and the cysteine variants in the isolated native outer membranes.

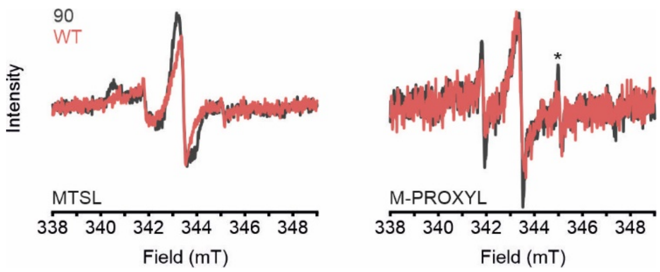

**Figure S4.** Room temperature continuous wave (cw) ESR spectra of selected nitroxide spin labels attached to BtuB wild-type (WT, red) or V90C (grey) in whole *E. coli* cells. The corresponding spin concentrations are given in Supplementary Table 1. Asterisk indicates a small amount of free spin label in the sample. As evident from the PELDOR data (see Figure S10), labeling failed for M-PROXYL at this position.

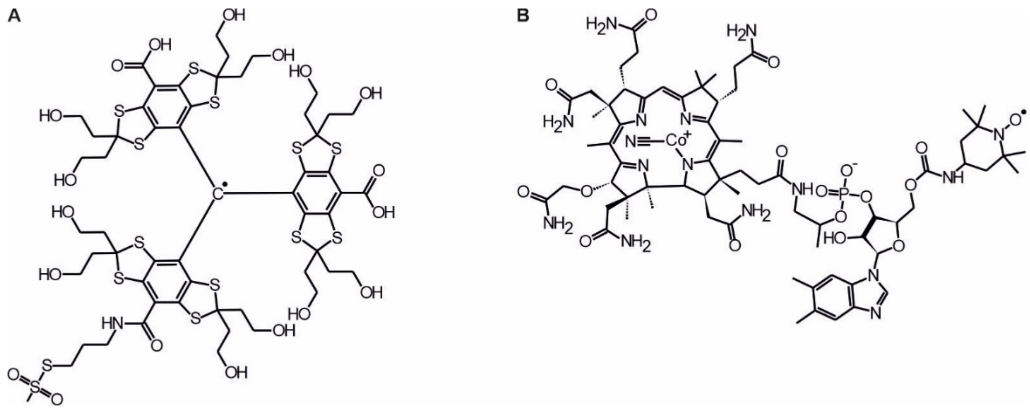

**Figure S5.** Chemical structure of (A) methanethiosulfonate-functionalized OX063 trityl spin label (MTS-OX063), and (B) of TEMPO-modified cyanocobalamin substrate (T-CNCbl).

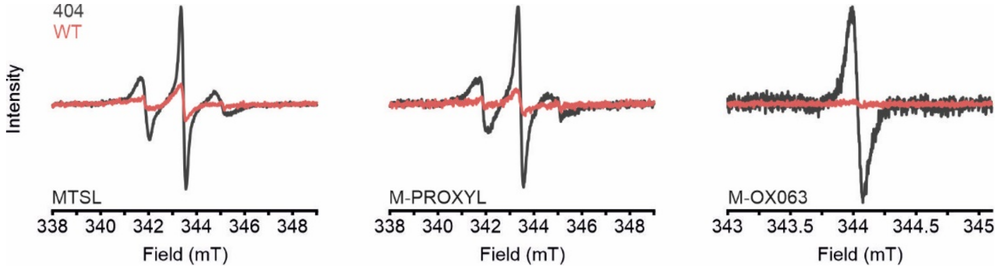

**Figure S6.** Room temperature cw ESR spectra of selected nitroxide and M-OX063 spin labels attached to BtuB wild-type (WT, red) or F404C (grey) in whole *E. coli* cells. The cw spectrum of BtuB F404C labeled with M-OX063 was measured using a smaller sample volume, which explains the lower S/N. The corresponding spin concentrations are given in Supplementary Table 1.

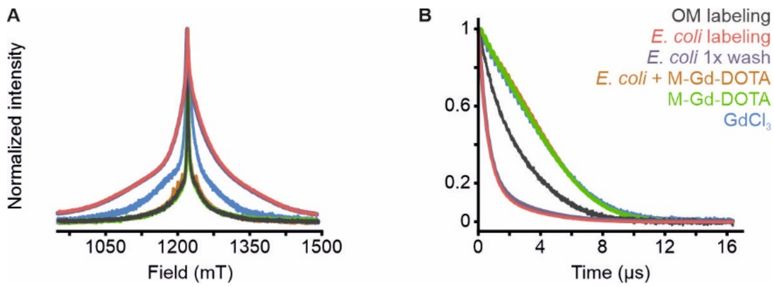

**Figure S7.** Pulsed ESR analysis of Gd(III) containing samples in *E. coli*, native outer membranes, and buffer. (A) Normalized echo-detected field-swept spectra recorded at 10 K, and (B) the corresponding transversal relaxation measurements. In contrast to native outer membranes, BtuB T188C could not be labeled using M-Gd-DOTA in *E. coli* and the PELDOR data did not give any distances (Figure S9C). The native outer membranes show a narrow central transition (in dark grey, $T_M$ = 2.6 µs) with a width comparable with free M-Gd-DOTA spin label (in green, $T_M$ = 4.5 µs). Labeling in whole cells lead to a significantly broadened spectrum (in red) and a reduced $T_M$ of 0.8 µs. Washing the cells 1x more before PELDOR sample preparation did not make any difference (in purple, underneath the red spectrum, $T_M$ = 0.9 µs). The spectrum of $GdCl_3$ solution ($T_M$ = 4.5 µs) is shown as a control (in blue). Direct addition of M-Gd-DOTA to *E. coli* followed by immediate sample freezing did not produce the broadening (in orange, $T_M$ = 4.7 µs). Altogether the data suggest a release of Gd(III) from the chelator during incubation of the cells with the spin label (at 15 µM). Fitting of the decay curves to obtain the $T_M$ values are shown in Figure S8E.

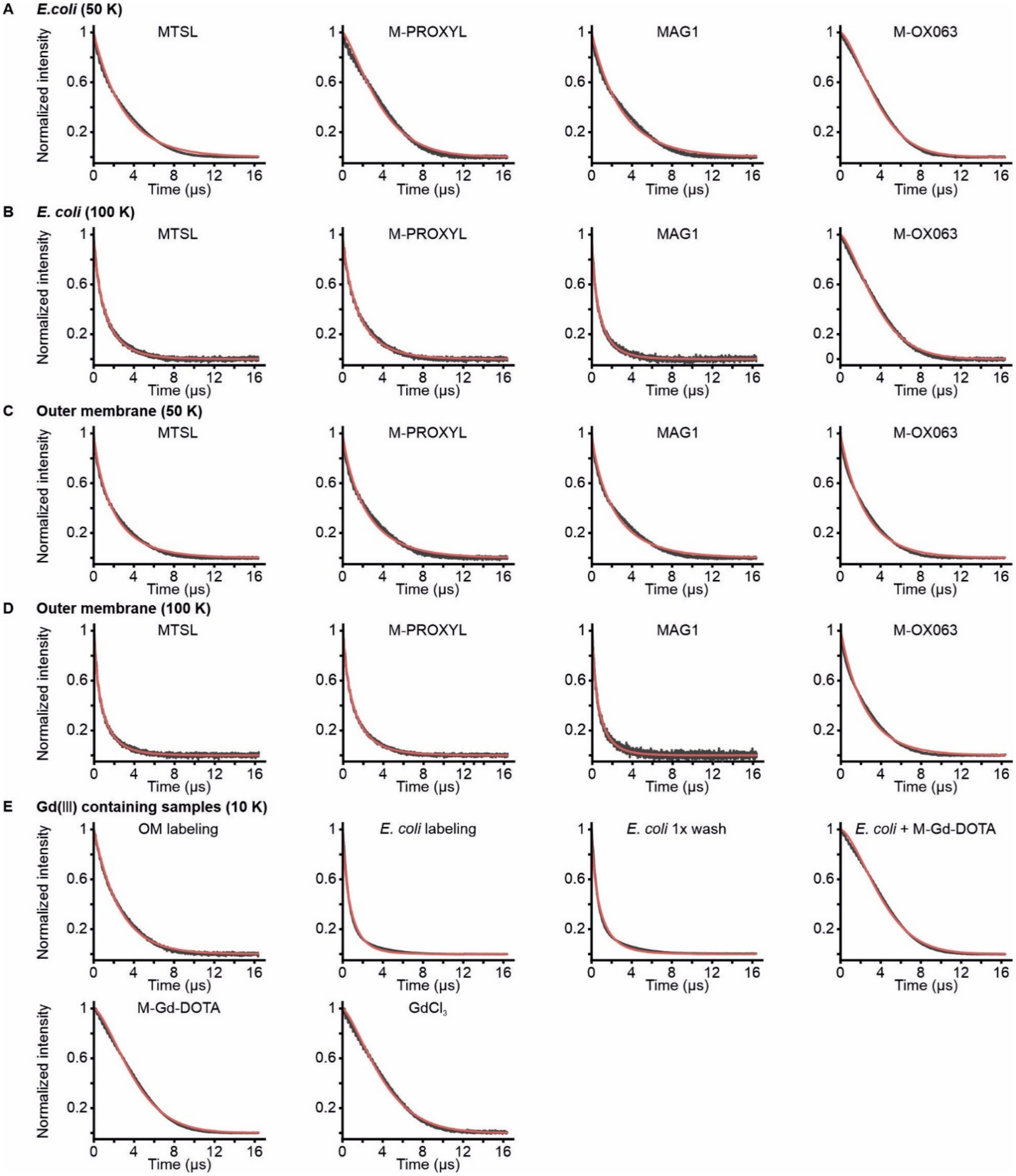

**Figure S8.** Fits with the stretched exponential function $\exp[-(2\tau/T_M)^{\kappa}]$ for the transversal relaxation curves for spin labeled BtuB T188C in *E. coli* and native outer membranes. (A) Fits (in red) to the original data (grey) obtained in *E. coli* cells at 50 K, or (B) at 100 K; (C) in native outer membranes at 50 K, or (D) at 100 K for the indicated spin labels. The values obtained for $T_M$ and $\kappa$ are summarized in Table 1. (E) Fits to the data of Gd(III) containing samples in *E. coli*, native outer membranes, and in buffer at 10 K as described in Figure S7.

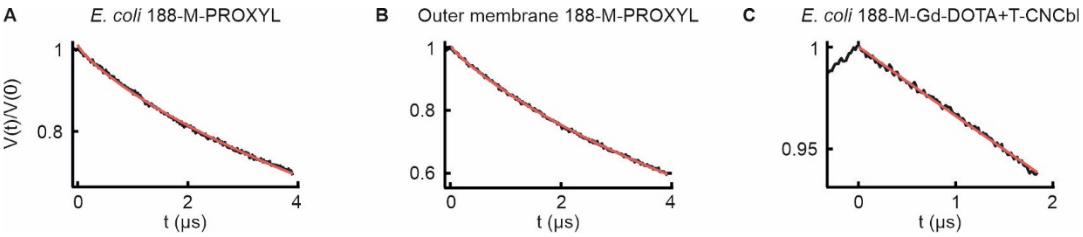

**Figure S9.** PELDOR spectroscopy of M-PROXYL or M-Gd-DOTA attached to BtuB T188C in *E. coli* and native outer membranes. (A) The primary PELDOR/DEER data of M-PROXYL labeled BtuB T188C (in the absence of T-CNCbl) in *E. coli* cells (A) and (B) in outer membranes at 50 K, which fit into stretched exponential decay functions (in red) with a dimensionality (*d*) of 2.40 and 2.64, respectively. (C) PELDOR data at 10 K for M-Gd-DOTA attached to BtuB T188C in whole *E. coli* cells in presence of T-CNCbl. The primary data fits into a mono-exponential decay (*d* = 3.0) devoid of any distances, revealing that M-Gd-DOTA labeling was not effective under the conditions employed.

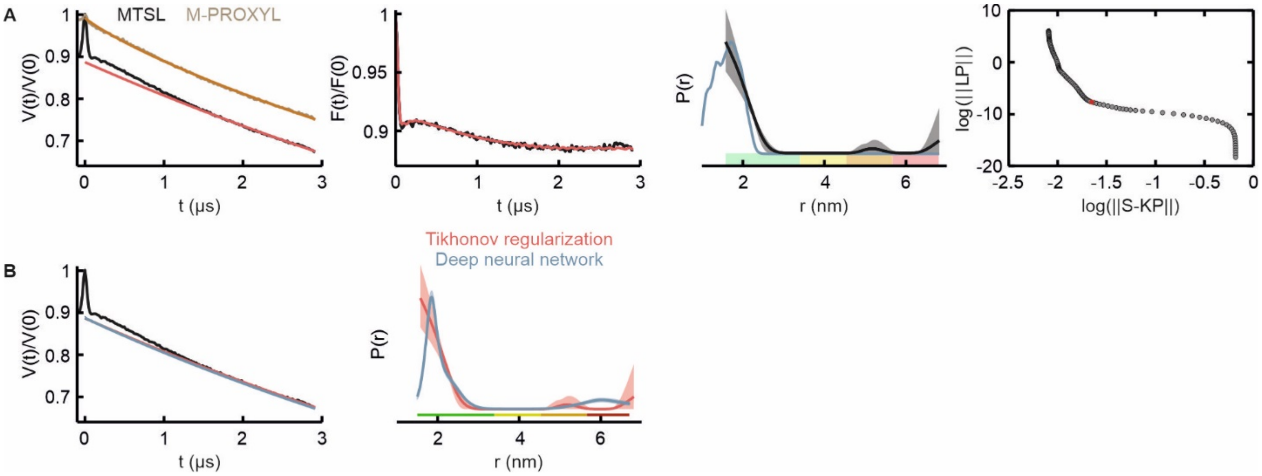

**Figure S10.** Data analysis of PELDOR/DEER measurements between indicated nitroxides attached to BtuB V90C and spin labeled substrate (T-CNCbl) in whole *E. coli* cells at 50 K. This position is located on the plug domain inside the barrel (Figure S2), overall, which has a limited accessibility. (A) Left panel shows the primary data (in grey for MTSL, in light brown for M-PROXYL) with the intermolecular (background) contribution overlaid (red for MTSL, orange for M-PROXYL). The data for M-PROXYL fits into a stretched exponential decay with a dimensionality of 2.6 and is devoid of any distances. Second panel, the background corrected form factor for MTSL overlaid with the corresponding fit. Third panel, the resulting distance distribution using Tikhonov regularization. The error bounds show variations of the probability distribution corresponding to the uncertainty in the background function (see Supplementary Table 2) and the reliability of shape, width, and mean distance are color coded as implemented in the DeerAnalysis software. The corresponding simulation on the crystal structure of BtuB is shown in light blue. Right most panel, the L-curve obtained from Tikhonov regularization with the chosen regularization point highlighted in red. (B) Left panel shows the primary data (in grey) overlaid with the intermolecular (background) contribution obtained from deep neural networks (blue) compared to manual Tikhonov regularization (red). Right panel, the obtained distance distributions with the error bounds.

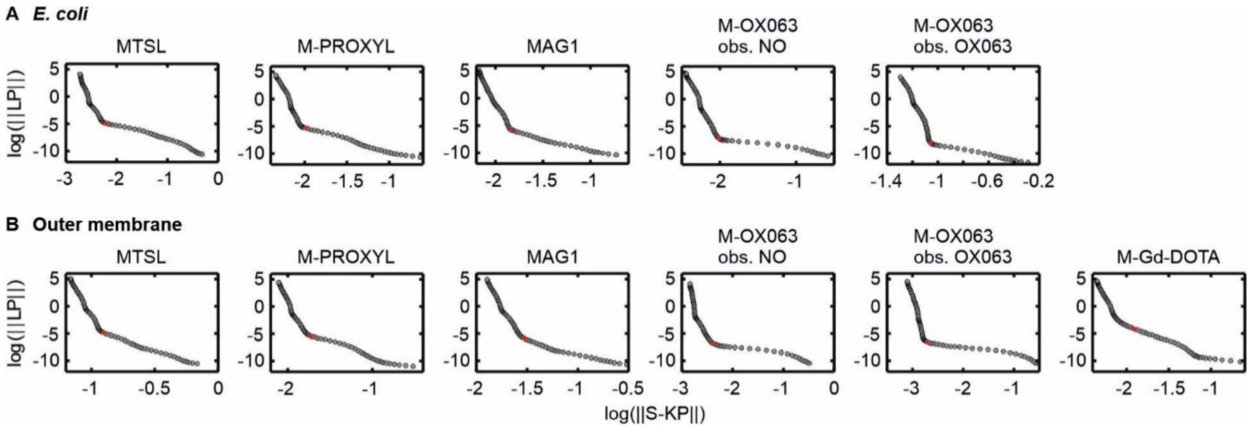

**Figure S11.** L-curves obtained from Tikhonov regularization for PELDOR measurements between BtuB T188C and the spin labeled substrate (T-CNCbl). (A) L-curves for nitroxide or M-OX063 labeled BtuB in whole *E. coli* cells corresponding to the data shown in Figure 6 with the used regularization parameter highlighted in red. (B) L-curves for the indicated labels in the native outer membranes corresponding to the data shown in Figure 7.

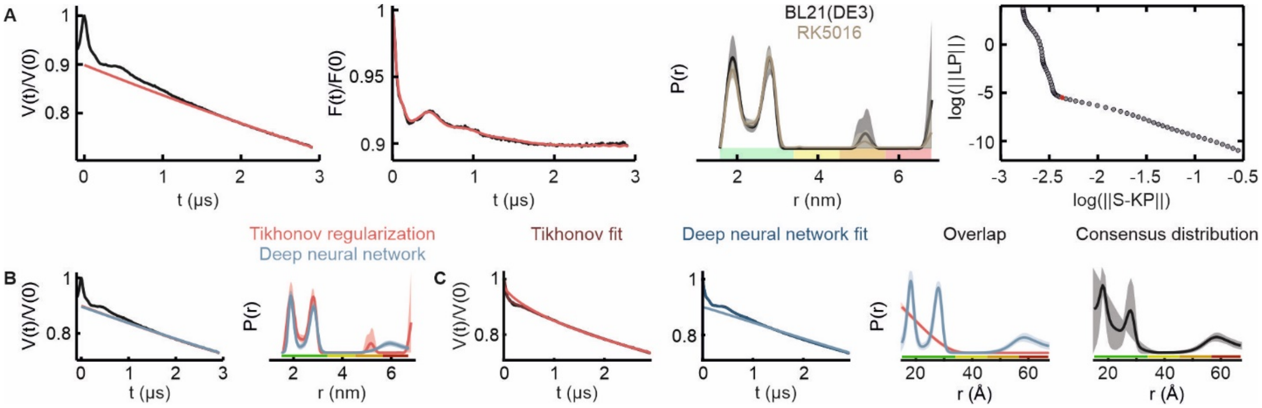

**Figure S12**. PELDOR data analysis for measurements between M-PROXYL labeled BtuB T188C and spin labeled substrate (T-CNCbl) in BL21(DE3) cells. (A) Left panel shows the primary data (in grey) with the intermolecular (background) contribution (red) overlaid; Second panel, the background corrected form factor overlaid with the corresponding fit; Third panel, the resulting distance distribution obtained using Tikhonov regularization overlaid with the probability distribution obtained in *E. coli* RK5016 cells (see Figure 6B); Right most panel with the L-curve obtained from Tikhonov regularization with the chosen regularization parameter highlighted in red. (B) Left panel shows the primary data (in grey) overlaid with the intermolecular (background) contribution obtained from deep neural network analysis (blue) compared to manual Tikhonov regularization (red); Right panel, the corresponding distance distributions and error bounds. (C) First and second panel present the experimental primary data (grey) overlaid with the fit and the intermolecular background contribution (in shades of red for Tikhonov, in shades of blue for deep neural networks) obtained using ComparativeDeerAnalyzer (CDA); Third panel, the corresponding distance distributions and error bounds; Right most panel, the resulting consensus distribution and uncertainty.

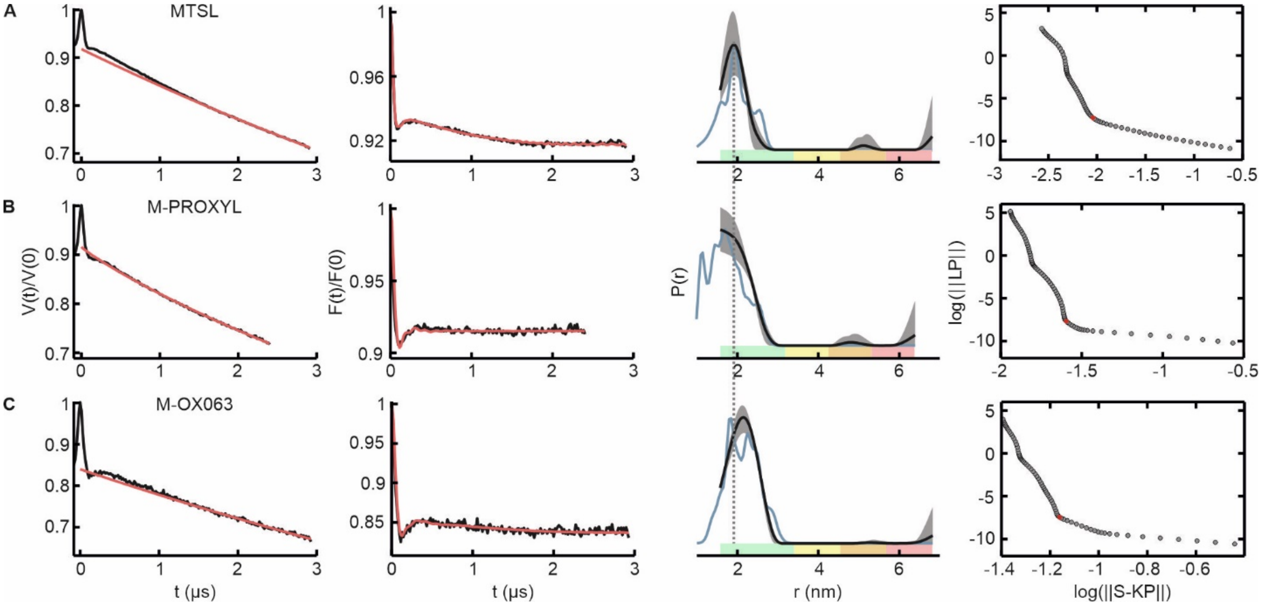

**Figure S13.** PELDOR data analysis for the indicated labels attached to BtuB F404C in whole *E. coli* cells. Distances were measured to the spin labeled substrate (T-CNCbl). (A-C) Left panels show the primary data (in grey) with the intermolecular (background) contribution overlaid (red); Second panels, the background corrected form factors overlaid with the corresponding fit; Third panels, the resulting distance distribution using Tikhonov regularization. Right panels, the L-curves obtained from Tikhonov regularization with the chosen regularization parameter highlighted in red. Measurements were performed at 50 K or 100 K while observing the nitroxide or the OX063, respectively. The corresponding simulations on the crystal structure of BtuB are overlaid in light blue.

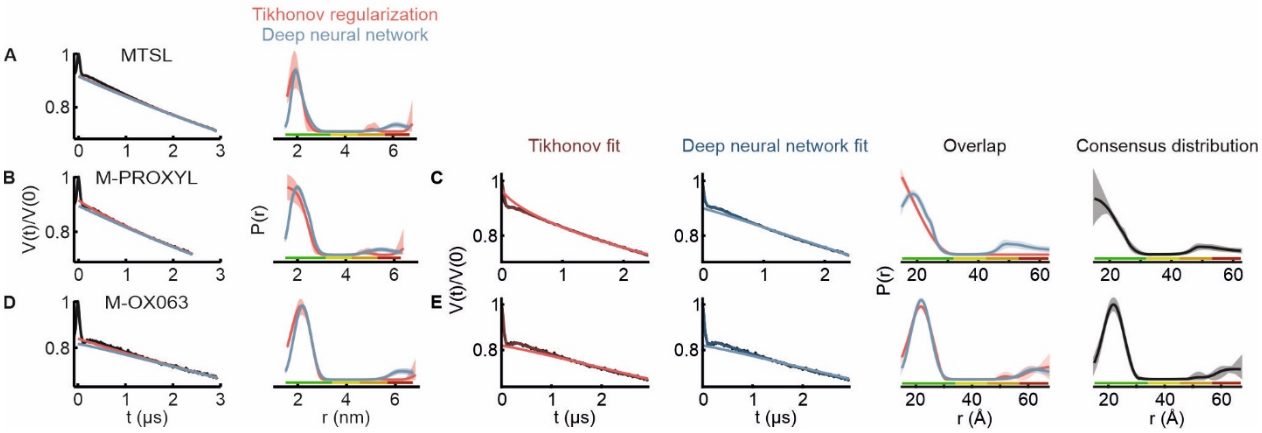

**Figure S14.** PELDOR data analysis for BtuB F404C labeled with the indicated labels in presence of spin labeled substrate (T-CNCbl) in whole *E. coli* cells using deep neural network (DEERNet) and ComparativeDeerAnalyzer (CDA). (A, B, D) Left panels show the primary data (grey) overlaid with the intermolecular (background) contribution obtained from deep neural networks (blue) compared to manual Tikhonov regularization (red) as presented in Figure S13; Right panels, the corresponding distance distributions and error bounds. (C, E) First and second panels present the experimental primary data (grey) overlaid with the fit and the intermolecular background contribution (in shades of red for Tikhonov, in shades of blue for deep neural networks) obtained using ComparativeDeerAnalyzer; Third panels, the corresponding distance distributions with the error bounds; Right panels, the resulting consensus distribution and uncertainty. For MTSL (A), ComparativeDeerAnalyzer output is not shown.

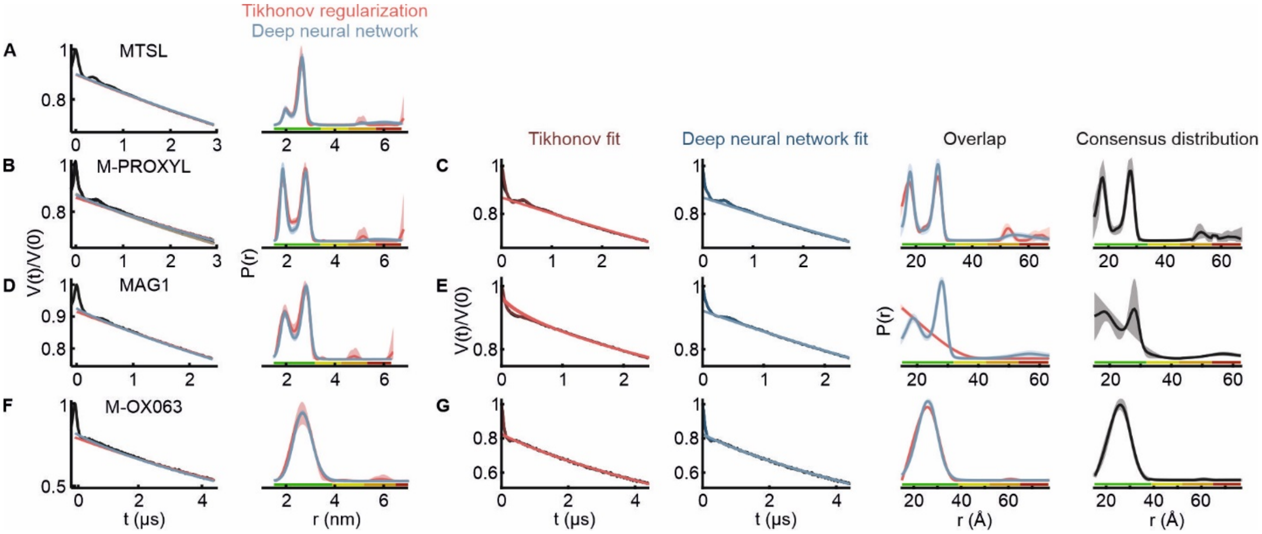

**Figure S15.** Data analysis of PELDOR measurements between BtuB T188C and spin labeled substrate (T-CNCbl) in whole *E. coli* cells using deep neural networks (DEERNet) and ComparativeDeerAnalyzer (CDA). (A, B, D, F) Left panels show the primary data (grey) overlaid with the intermolecular (background) contribution obtained from deep neural networks (blue) compared to manual Tikhonov regularization (red) as presented in Figure 6; Right panels, the corresponding distance distributions with the error bounds. (C, E, G) First and second panels present the experimental primary data (grey) overlaid with the fit and the intermolecular background contribution (in shades of red for Tikhonov, in shades of blue for deep neural networks) obtained using ComparativeDeerAnalyzer; Third panels, the corresponding distance distributions and error bounds; Right most panels, the resulting consensus distribution and the uncertainty. For MTSL, ComparativeDeerAnalyzer output is not shown.

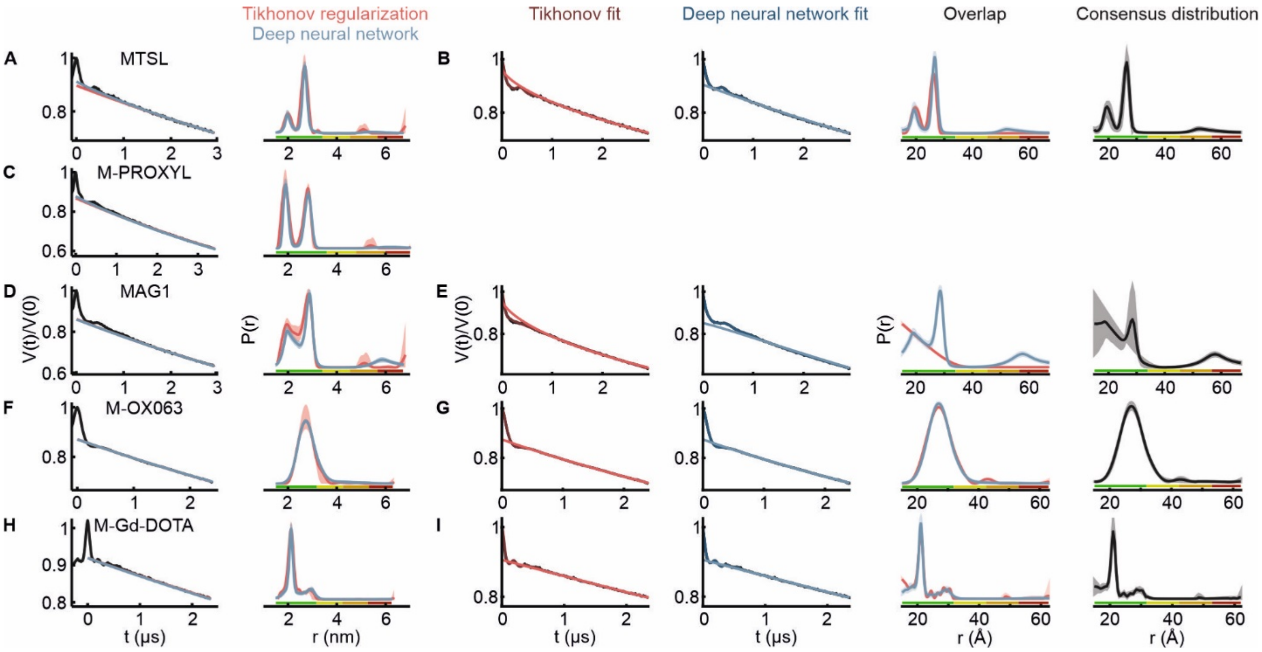

**Figure S16.** Data analysis of PELDOR/DEER measurements between BtuB T188C and spin labeled substrate (T-CNCbl) in native outer membranes at 50 K using deep neural networks (DEERNet) and ComparativeDeerAnalyzer (CDA). (A, C, D, F, H) Left panels show the primary data (grey) overlaid with the intermolecular (background) contribution obtained from deep neural networks (blue) compared to manual Tikhonov regularization (red) as presented in Figure 7; Right panels, the corresponding distance distributions with the error bounds. (B, E, G, I) First and second panels present the primary data (grey) overlaid with the fit and the intermolecular background contribution (in shades of red for Tikhonov, in shades of blue for deep neural network analysis) obtained using ComparativeDeerAnalyzer; Third panels, the corresponding distance distributions with the error bounds; Right most panels, the resulting consensus distribution with the uncertainty. For M-PROXYL, ComparativeDeerAnalyzer output is not shown.

### Characterization of M-OX063

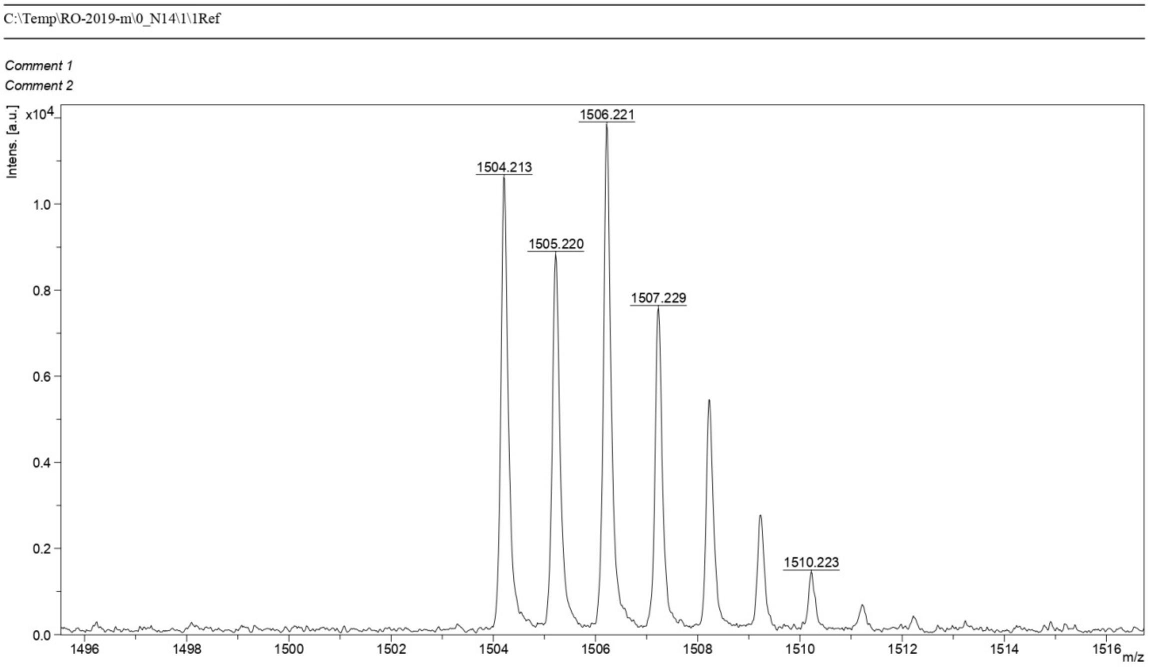

**Figure S17.** HRMS MALDI-TOF spectrum of trityl M-OX063. Calculated m/z for $C_{58}H_{69}N_2NaO_{19}S_{12}$ $[M^{+}Na]^{+}$ is 1504.1041; observed m/z is 1504.213.

| | | | | | | | |
|---|---|---|---|---|---|---|---|
| *Title* | RO-2000-3 KBr 150:1 Tensor 27 | | | *Origin* | gmkk | *File Name* | D:\MY PERSONAL\TMP\IR ARCHIVE\2019\GMKKRO-2000-3.DX |
| *Date Stamp* | 22/8/2019 | *Date* | 22 Aug 2019 14:00:06 | | | *Technique* | Infrared; *Spectral Region* IR |
| *X Axis* | Wavenumber (cm-1) | | | *Y Axis* | Transmittance | *Spectrum Range* | 399.2111 - 3997.8959 |
| *Points Count* | 1867 | *Data Spacing* | 1.9286 | | | | |

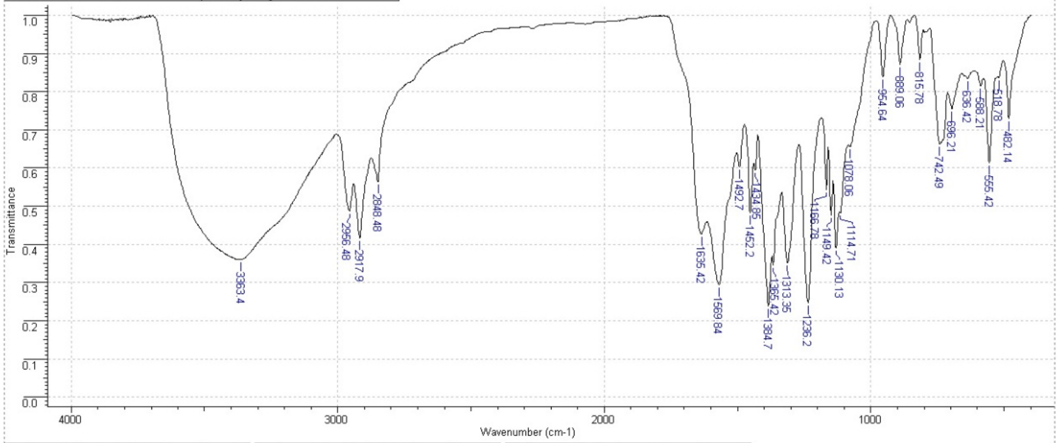

| No | cm-1 | T | FWHH | Asym | Intensity |
|---|---|---|---|---|---|
| 1 | 3363.40 | 0.361 | -1.00 | 0.00 | S |
| 2 | 2956.48 | 0.488 | - | - | S |
| 3 | 2917.90 | 0.417 | - | - | S |
| 4 | 2848.48 | 0.564 | - | - | M |
| 5 | 1635.42 | 0.428 | - | - | S |
| 6 | 1569.84 | 0.295 | - | - | VS |
| 7 | 1492.70 | 0.603 | - | - | M |
| 8 | 1452.20 | 0.483 | - | - | S |
| 9 | 1434.85 | 0.595 | - | - | M |
| 10 | 1384.70 | 0.239 | - | - | VS |

| No | cm-1 | T | FWHH | Asym | Intensity |
|---|---|---|---|---|---|
| 11 | 1365.42 | 0.346 | - | - | S |
| 12 | 1313.35 | 0.351 | - | - | S |
| 13 | 1236.20 | 0.248 | -1.00 | 0.00 | VS |
| 14 | 1166.78 | 0.544 | - | - | M |
| 15 | 1149.42 | 0.478 | - | - | S |
| 16 | 1130.13 | 0.391 | - | - | S |
| 17 | 1114.71 | 0.484 | - | - | S |
| 18 | 1078.06 | 0.657 | - | - | M |
| 19 | 954.64 | 0.839 | -1.00 | 0.00 | W |
| 20 | 889.06 | 0.872 | -1.00 | 0.00 | W |

| No | cm-1 | T | FWHH | Asym | Intensity |
|---|---|---|---|---|---|
| 21 | 815.78 | 0.884 | -1.00 | 0.00 | W |
| 22 | 742.49 | 0.663 | - | - | M |
| 23 | 696.21 | 0.755 | - | - | M |
| 24 | 636.42 | 0.834 | - | - | W |
| 25 | 588.21 | 0.817 | - | - | W |
| 26 | 555.42 | 0.614 | -1.00 | 0.00 | M |
| 27 | 518.78 | 0.838 | - | - | W |
| 28 | 482.14 | 0.729 | -1.00 | 0.00 | M |

**Figure S18.** IR spectrum (KBr) of trityl M-OX063 (KBr). Further details are given in the methods section.

### Supplementary Tables

**Supplementary Table 1.** Spin concentration for nitroxide spin labels and M-OX063 in *E. coli* cells and native outer membranes overexpressing BtuB wild-type (WT) or the cysteine variants V90C, T188C, and F404C. The given values for outer membrane samples correspond to the total spin concentration of the membrane containing ~15 µM of BtuB. For *E. coli* samples, the spin concentration is normalized to the same $OD_{600}$ value. A maximum of 20 % error is estimated for the spin concentration. Values in brackets possess a larger uncertainty due to poor signal to noise. For *E. coli* cells overexpressing BtuB WT labeled with M-OX063, the spin concentration was below the detectable limit (denoted as b.d.).

| Spin label | ***E. coli*** **(µM)** | | | | **Outer membrane (µM)** | |
|---|---|---|---|---|---|---|
| | **WT** | **V90C** | **T188C** | **F404C** | **WT** | **T188C** |
| MTSL | (15±3) | 21±4 | 26±5 | 27±5 | 59±12 | 85±17 |
| M-PROXYL | (8±2) | (9±2) | 21±4 | 22±4 | 40±8 | 77±15 |
| MAG1 | (13±3) | - | 29±6 | - | 67±13 | 75±15 |
| M-OX063 | b.d. | - | 4±1 | 4±1 | 10±2 | 26±5 |

**Supplementary Table 2.** Error estimation for PELDOR/DEER measurements between cysteine variants of BtuB and spin labeled substrate (T-CNCbl) in *E. coli* cells and native outer membranes. The data were analyzed using Tikhonov regularization as featured in the DeerAnalysis2018 software package. Probability distributions were calculated for different background models from a combined variation of the starting time (in the indicated range in 11 steps) and the dimensionality of the background function (*d* = 2.4–3.0 in 3 steps). The range values for *d* were defined based on the measurement of single cysteine variants (see Figure S9). A prune level $L_{\text{prune}}$ of 1.15 of the root mean square deviation (r.m.s.d.) of the best fit was used.

| Sample | Figure(s) | Error estimation/validation | | | | | Regularization parameter (α) |
|---|---|---|---|---|---|---|---|
| | | Dimensionality (d) | | Starting time window | | | |
| | | Value/range | Steps | $t_{max}$ (μs) | Range (ns) | Steps | |
| **In *E. coli* cells** | | | | | | | |
| 188-MTSL (RK5016) | 6, S15 | 2.4-3 | 3 | 2.93 | 576-1744 | 11 | 7.94 |
| 188-M-PROXYL (RK5016) | 6, S15 | 2.4-3 | 3 | 2.91 | 576-1744 | 11 | 12.6 |
| 188-M-PROXYL (BL21(DE3)) | S12 | 2.4-3 | 3 | 2.93 | 576-1744 | 11 | 12.6 |
| 188-MAG1 (RK5016) | 6, S15 | 2.4-3 | 3 | 2.41 | 480-1440 | 11 | 15.8 |
| 188-M-OX063 (RK5016) | 6, S15 | | | | | | |
| Obs. OX063 | | 2.4-3 | 3 | 4.38 | 880-2624 | 11 | 316 |
| Obs. NO | | 2.4-3 | 3 | 2.38 | 480-1440 | 11 | 63.1 |
| 404-MTSL (RK5016) | S13, S14 | 2.4-3 | 3 | 2.93 | 576-1744 | 11 | 50.1 |
| 404-M-PROXYL (RK5016) | S13, S14 | 2.4-3 | 3 | 2.41 | 480-1440 | 11 | 100 |
| 404-M-OX063 (RK5016) obs. NO | S13, S14 | 2.4-3 | 3 | 2.93 | 592-1760 | 11 | 126 |
| 90-MTSL (RK5016) | S10 | 2.4-3 | 3 | 2.91 | 576-1744 | 11 | 158 |
| **In the native outer membrane (OM)** | | | | | | | |
| 188-MTSL (RK5016) | 7, S16 | 2.4-3 | 3 | 2.91 | 576 – 1744 | 11 | 15.9 |
| 188-M-PROXYL (RK5016) | 7, S16 | 2.4-3 | 3 | 3.42 | 688-2048 | 11 | 15.9 |
| 188-MAG1 (RK5016) | 7, S16 | 2.4-3 | 3 | 2.93 | 576 – 1744 | 11 | 25.1 |
| 188-M-OX063 (RK5016) | 7, S16 | | | | | | |
| Obs. OX063 | | 2.4-3 | 3 | 2.35 | 480-1440 | 11 | 25.1 |
| Obs. NO | | 2.4-3 | 3 | 2.40 | 480-1456 | 11 | 39.8 |
| 188-M-Gd-DOTA (RK5016) | 7, S16 | 2.4-3 | 3 | 2.35 | 464-1408 | 11 | 3.98 |
